# Chromogranin A (CgA) Deficiency Attenuates Tauopathy by Altering Epinephrine–Alpha-Adrenergic Receptor Signaling

**DOI:** 10.1101/2024.06.11.598548

**Authors:** Suborno Jati, Daniel Munoz-Mayorga, Shandy Shahabi, Kechun Tang, Yuren Tao, Dennis W. Dickson, Irene Litvan, Gourisankar Ghosh, Sushil K. Mahata, Xu Chen

## Abstract

Our previous studies have indicated that insulin resistance, hyperglycemia, and hypertension in aged wild-type (WT) mice can be reversed in mice lacking chromogranin-A (CgA-KO mice). These health conditions are associated with a higher risk of Alzheimer’s disease (AD). CgA, a neuroendocrine secretory protein has been detected in protein aggregates in the brains of AD patients. Here, we determined the role of CgA in tauopathies, including AD (secondary tauopathy) and corticobasal degeneration (CBD, primary tauopathy). We found elevated levels of CgA in both AD and CBD brains, which were positively correlated with increased phosphorylated tau in the frontal cortex. Furthermore, CgA ablation in a human P301S tau (hTau) transgenic mice (CgA-KO/hTau) exhibited reduced tau aggregation, resistance to tau spreading, and an extended lifespan, coupled with improved cognitive function. Transcriptomic analysis of mice cortices highlighted altered levels of alpha-adrenergic receptors (Adra) in hTau mice compared to WT mice, akin to AD patients. Since CgA regulates the release of the Adra ligands epinephrine (EPI) and norepinephrine (NE), we determined their levels and found elevated EPI levels in the cortices of hTau mice, AD and CBD patients. CgA-KO/hTau mice exhibited reversal of EPI levels in the cortex and the expression of several affected genes, including Adra1 and 2, nearly returning them to WT levels. Treatment of hippocampal slice cultures with EPI or an Adra1 agonist intensified, while an Adra1 antagonist inhibited, tau hyperphosphorylation and aggregation. These findings reveal a critical role of CgA in regulation of tau pathogenesis via the EPI-Adra signaling axis.

## Introduction

Tauopathies are a broad class of neurodegenerative diseases characterized by the accumulation of tau aggregates in the brain. Primary tauopathies such as corticobasal degeneration (CBD) and Pick’s disease (PiD) exhibit only tau inclusions, whereas Alzheimer’s disease (AD), a secondary tauopathy, is characterized by the presence of extracellular beta-amyloid (Aβ) plaques and intracellular neurofibrillary tau tangles (*1–4*). Hyperphosphorylation of tau weakens tau’s microtubule-binding affinity, leading to its neurotoxic aggregation, forming tangles of different substructures (*5*). Tauopathies are closely associated with metabolic dysfunctions, including insulin resistance, hyperglycemia, and hypertension (*6, 7*), as well as neuroinflammation (*8, 9*). However, the pathophysiology underlying tauopathy development during aging remains elusive.

CgA, along with its paralog CgB, co-stores catecholamines, calcium, and neuropeptides in dense-core vesicles (DCVs) present in cells of the neuroendocrine system and regulates the secretion of the vesicular content from the cell (*10, 11*). Depletion of CgA in WT mice improves their systemic glucose metabolism and hypertension compared to WT mice, suggesting that CgA is an aging risk factor (*12*). Incidentally, aging is the greatest risk factor for sporadic AD (sAD), which represents nearly 95% of all AD cases (*13*). CgA was reported to be present in proteinopathic aggregates (*14, 15*) and is elevated in AD brain (*16*). As a prohormone generating various bioactive peptides, CgA and its peptides impact age-related diseases such as hypertension and diabetes (*17–19*). CgA is also known to induce chronic inflammation (*20*), suggesting its potential role in facilitating primary and secondary tauopathies, including AD .

In this study, we first determined the changes of CgA levels and how they correlate with tau pathology, in the cortices of AD, corticobasal degeneration (CBD), and hTau mice, where the human Tau (1N4R) protein with the P301S mutation is overexpressed under the PRNP (prion protein gene) promoter (*21*). Subsequently, we explored the effects of CgA ablation in hTau mice with regard to neuropathology, cognitive function and transcriptomics. Because CgA is known to play a pivotal role in the adrenergic system, we explored the alpha-adrenergic receptor 1 (Adra1) and epinephrine (EPI) signaling axis in hTau mice, and the inflammatory response profile. Using organotypic hippocampus slice cultures, we tested the hypothesis that CgA regulates tau pathogenesis via activating EPI-Adra1 signaling and promoting inflammation. Our results suggest that the EPI-Adra1 system plays a crucial role in tau pathogenesis, which is modulated by CgA, highlighting a novel mechanistic connection between metabolic regulation and neurodegeneration.

## Results

### CgA level is elevated in the brains of AD, CBD and hTau mice and is associated with Tau tangles

We tested the levels of CgA in AD and CBD patients to find any correlation between disease pathology and CgA protein levels. Western blot (WB) analysis revealed a statistically significant increase in CgA protein levels in Braak stage VI (advanced stage) patient frontal cortex lysates, compared to Braak stage I/II (early stage) (**Fig 1A–B**). Elevated CgA in Braak VI samples positively correlated with higher levels of pathogenic tau phosphorylation (**Supplementary Fig 1A**). Immunohistochemical (IHC) analyses on the hippocampi of AD patients using a CgA-specific antibody showed increased CgA accumulation in Braak VI samples compared to Braak I/II samples (**Fig 1D–E**). CgA mRNA levels, however, remained unchanged across different Braak stages, suggesting that CgA protein stabilization occurs with the disease (**Fig 1C**). Co-staining of hippocampi from Braak VI samples revealed that CgA and tau aggregates co-reside, indicating that CgA might be associated with tau aggregates (**Supplementary Fig 1E**). In contrast, minimal overlap was observed in a Braak I hippocampus sample (**Supplementary Fig 1F**).

**Figure 1.**
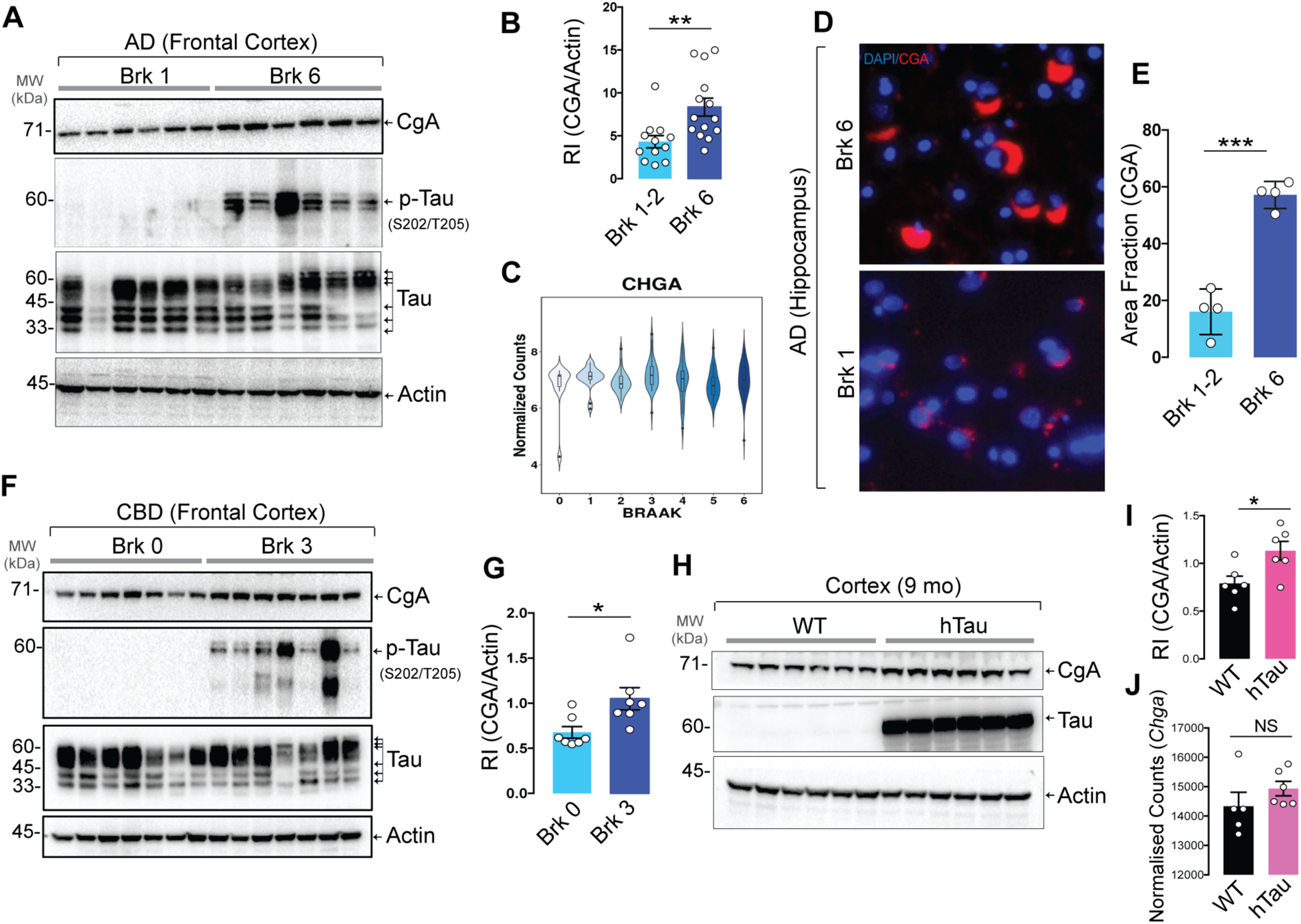
Augmented CgA protein levels in AD and CBD patient samples and P301S hTau transgenic mice (PS19). A. Representative Western blot (WB) of postmortem frontal cortex extracts showing levels of CgA, p-Tau, total Tau and Actin in six Braak stage 6 (Brk 6) and Braak stage 1/2 (Brk 1/2) postmortem patient frontal cortex lysates. B. Quantification of WB in (A) and other samples (Brk 0-2, n=12; Brk6, n=14). C. Quantification of *CHGA* (gene encoding CgA) transcript levels of patient samples from all six Braak stages. The RNAseq data was obtained from publicly available AD-AMP dataset. D. Representative immunofluorescence (IF) images showing CgA signalin the hippocampus of Brk 6 and Brk 1-2 samples. E. Image J quantification of the immunoreactive CgA in the hippocampus shown as area fraction (%) of total hippocampus (n=4 per group). F. WB of frontal cortex lysates of CBD and normal control patients showing levels of CgA, pTau, Tau and actin. G. Quantification of CgA levels in WB shown in F, normalized to actin levels (n=7 per group). H. WB of frontal cortex lysates of WT and hTau mice for CgA, Tau and actin. I. Quantification of WB shown in H for CgA levels, normalized to actin levels (n=6 per group). J. Transcript level of CgA in WT (n=5) and hTau mice (n=6). Values are average +/-SEM. p-values were calculated using unpaired two-tailed T-test. \**p* < 0.05, \*\**p* < 0.01, \*\*\**p* < 0.001, \*\*\*\**p* < 0.0001

In addition to AD, we also found elevated CgA levels in CBD-affected frontal cortices compared to normal controls (**Fig 1F–G**). Similarly, CgA levels were positively correlated with phospho-tau levels (**Supplementary Fig 1B**). Consistent with human samples, WB showed higher CgA protein levels in the cortices of hTau mice compared to WT mice (**Fig 1H–I**) without any change in the transcript levels (**Fig 1J**). We further confirmed higher levels of CgA in the CA3 region of the hippocampus through IHC (**Supplementary Fig 1C–D**).

To determine how CgA depletion per se affects tau phosphorylation and aggregation, we generated hippocampal organotypic slice cultures (OTSC) from WT and CgA-KO pups (7–8 days old) followed by AAV-mediated transduction of TauP301S and seeding with synthetic tau fibrils (K18/PL, (*22*)) (**Supplementary Fig 1G**). Compared to CgA-KO slices, WT slices had more pronounced pathological tau phosphorylation (**Supplementary Fig 1H–J**), as well as tau aggregation, as revealed by a tau-aggregation-specific monoclonal antibody (MC1; P < 0.05; **Supplementary Fig 1K–L**). These observations indicate reduced tau phosphorylation and aggregation upon CgA depletion.

### Genetic deletion of CgA reduces neurodegeneration in tauopathy mice

To gain deeper insights into the role of CgA in tau-induced pathogenesis, we bred hTau mice with CgA-KO mice to generate CgA-KO/hTau mice. At nine months of age, Nissl staining revealed clear brain atrophy with reduced hippocampal volume in hTau mice, consistent with the previous report (*21*) . In contrast, CgA-KO/hTau mice exhibited minimal brain atrophy (**Fig 2A–B**; **Supplementary Fig 2A–B**). Corroborating these findings, WB analysis demonstrated reduced pathogenic tau hyperphosphorylation at S202/T205 (AT8) and S396/S404 (PHF1) residues in the cortex of CgA-KO/hTau mice compared to hTau mice (**Fig 2C**; **Supplementary Fig 2C–D**). We also observed reduced AT8 immunostaining in hippocampi of CgA-KO/hTau mice (**Supplementary Fig 2F–G**), corroborating our immunoblot results with cortices (**Fig 2C**). Furthermore, immunostaining with the MC1 antibody revealed significantly higher tau aggregation in the hippocampal CA3 cell-body region and granular cell layer of the dentate gyrus in nine-month-old hTau mice compared to CgA-KO/hTau mice (**Fig 2D–F**). These results suggest that CgA promotes tau phosphorylation and aggregation.

**Figure 2.**
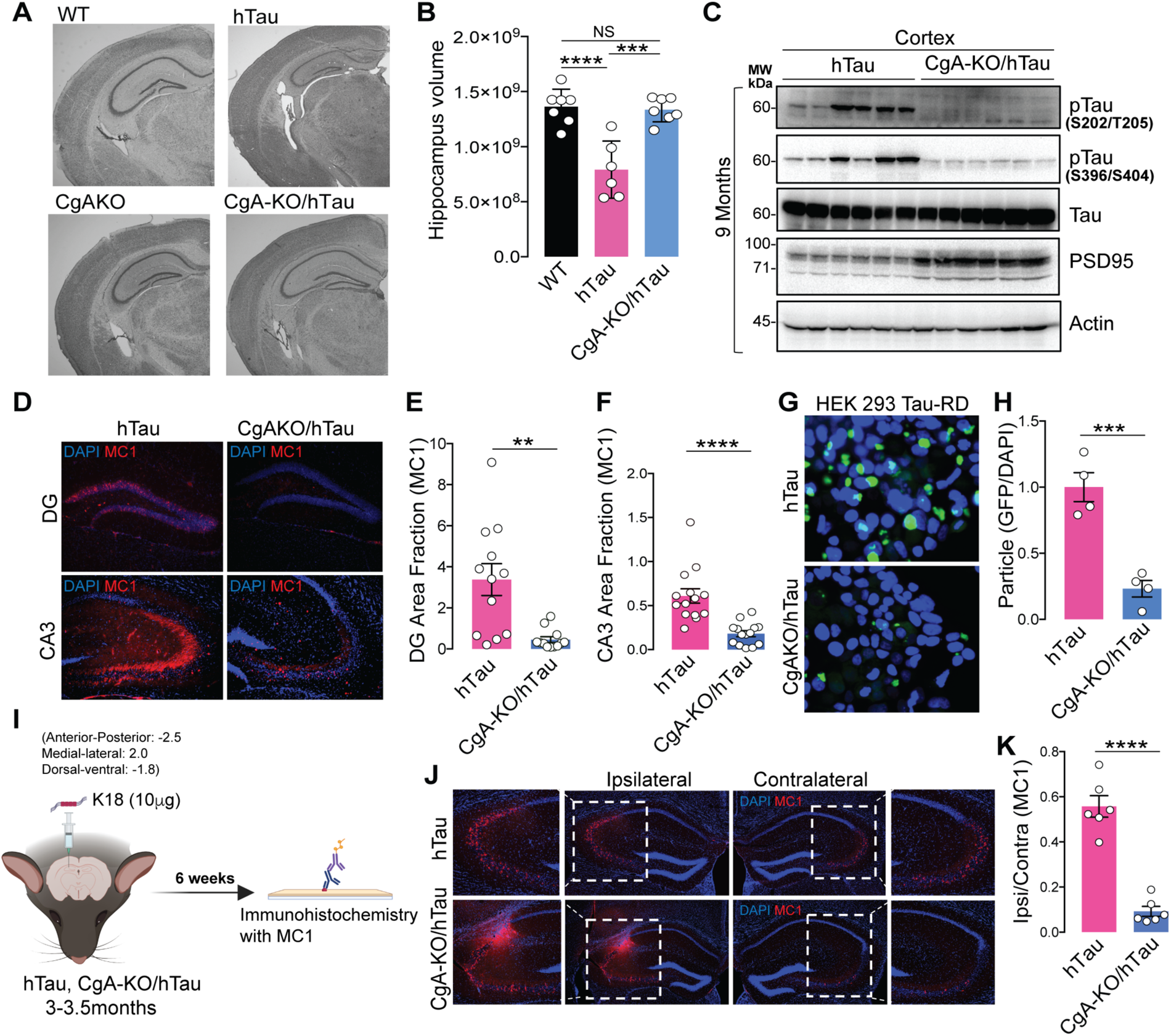
CgA deficiency reduces tauopathy. A. Representative Nissl staining of mouse brain section of WT, hTau, CgA-KO and CgA-KO/hTau littermate mice of 9-10 months of age. B. Estimated volume of the hippocampus of 9-10-months-old WT (n=7), hTau (n=6) and CgA-KO/hTau mice (n=7). C. WB of cortex RIPA lysates of hTau (n=6) and CgA-KO/hTau (n=6) mice showing levels of pathogenic phosphorylated Tau (pTau), total Tau, PSD95 and actin. D. Representative IF images showing the levels of MC1 labeled tau aggregates in the Dentate Gyrate (DG) and CA3 region of hippocampus. E. Image J quantification of MC1+ Tau aggregates revealed by IF in DG (n=10). F. Image J quantification of MC1+ Tau aggregates revealed by IF in CA3 (n=12). G. Fluorescence images of HEK293 cells expressing TauRD-GFP transfected with the brain extracts of hTau (top) or CgA-KO/hTau mice. H. Image J quantification of Tau aggregates (FRET+) as shown in F, normalized to DAPI counts (n=4 wells per group). I. Schematics describing the Tau fibrils (K18/PL)-induced spreading assay. J. Representative IF images using MC1 antibody showing spreading of tau aggregates from the site of injection (ipsilateral) to the opposite site (contralateral) of the hippocampus. A zoomed-in view of the aggregate-rich region is shown in each case. K. Image J quantification of tau aggregates as a ratio of MC1+ area in ipsilateral hippocampus relative to the contralateral hippocampus as in (J) (n = 6 mice per group). Values are average +/-SEM. p-values in B was calculated using one-way ANOVA followed by Tukey’s Multiple Comparison test. p-values in E, G and J was calculated using unpaired two-tailed T-test. \**p* < 0.05, \*\**p* < 0.01, \*\*\**p* < 0.001, \*\*\*\**p* < 0.0001

To quantify the difference in the level of seeding-competent tau aggregates, we delved deep into CgA’s role in tau aggregation and employed the well-established Förster resonance energy transfer (FRET) assay in HEK 293T cells expressing tau fused independently to GFP and RFP (*23*). The FRET signal measured in crude brain lysate was found to be higher in hTau mice than in CgA-KO/hTau mice, indicating lower tau seeding capacity in the brain lysates of CgA-KO/hTau mice compared to hTau mice (**Fig 2G–H**). We further investigated CgA’s impact on tau seeding and spreading by stereotaxically injecting synthetic K18 tau fibrils into the dentate gyrus (DG) of three-month-old hTau and CgA-KO/hTau mice (**Fig 2I**). Fibril-induced tau spread to the CA3 region of the contralateral hippocampus was evident in hTau mice 6 weeks post-injection, but minimal in CgA-KO/hTau mice, suggesting that CgA promotes tau spread (**Fig 2J-K**).

We further characterized neurodegeneration by WB with an anti-PSD95 antibody and by ultrastructural analysis via electron microscopy (EM) and found increased PSD95 levels in the cortex (**Fig 2C**; **Supplementary Fig 2E**) and amplified synapse numbers (marked by arrows) in the hippocampus of CgA-KO/hTau mice compared to hTau mice (**Supplementary Fig 2E, 2J– K**). Notably, hTau mice had fewer synaptic vesicles (SVs) in their presynaptic neurons and irregular synapses compared to CgA-KO/hTau mice (**Supplementary Fig 2L–M**). Collectively, these findings suggest that CgA plays a pro-neurodegenerative role in tau-induced pathogenesis.

### CgA deficiency improves learning, memory, cognition, and longevity of hTau mice

We conducted behavioral assessments on four distinct genotypes of mouse littermates (WT, CgA-KO, hTau, and CgA-KO/hTau). In the Morris water maze (MWM) test, during the training phase, hTau mice traveled prolonged distances and durations before reaching the platform, and exhibited lack of improvement over the seven-day training period, compared to WT and CgA-KO mice (**Fig 3A–C**; **Supplementary Fig 3A**), indicating impaired spatial learning. In contrast, CgA-KO/hTau mice exhibited a complete rescue of the learning deficit (**Fig 3A–C**; **Supplementary Fig 3A**). The average latency in the probe trial for hTau mice was significantly longer than WT mice, indicative of memory impairment, whereas CgA-KO/hTau mice performed indistinguishably from the WT, spending significantly less time and more entries to the target zone during the probe trial (**Fig 3D, E**). Notably, both body weight and swimming speed remained comparable between hTau and CgA-KO/hTau mice (**Supplementary Fig 3B**; **Fig 3F**), indicating that the reduced latency in CgA-KO/hTau mice was not due to increased speed. In the novel object recognition (NOR) test, CgA-

**Figure 3.**
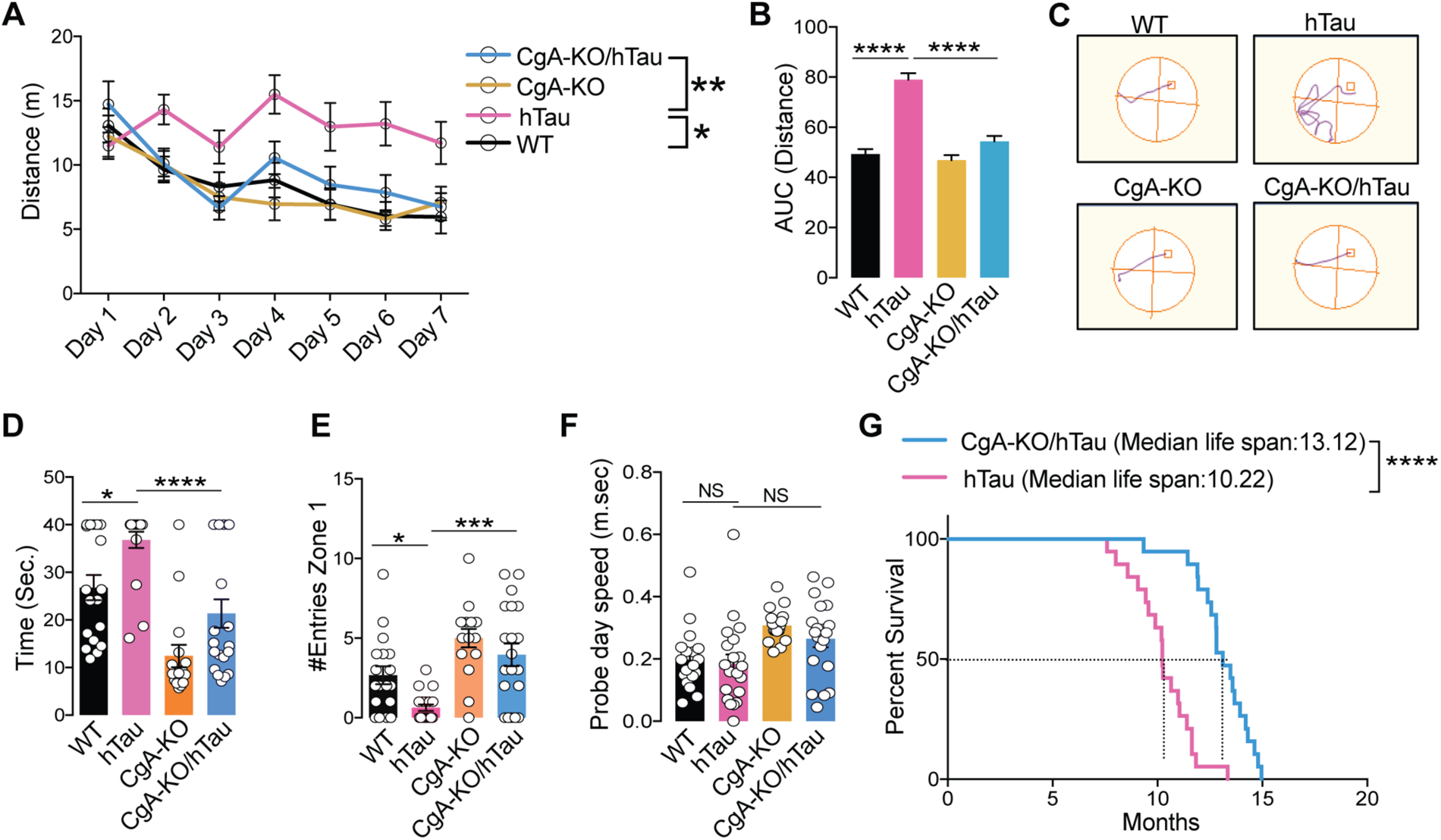
Ablation of CgA improved spatial learning and memory, and extended the life span of hTau mice. A. Distance travelled each day to reach the platform during seven days training trials of the Morris Water-Maze Test. CgA-KO/hTau (n=20), CgA-KO (n=16), hTau (n=19) and WT (n=18). B. Area Under the Curve (AUC) for the distance travelled during the training period. C. Probe trial performed 24 hr after the 7^th^ day of training for all the mice. The swimming paths in the probe trial of representative mice of all four groups are shown (the square marks the location of the platform during the goal acquisition trials). D. Time taken (latency) to reach the target platform. E. Number of entries in the zone where the platform was present (correct zone). F. Swimming speed of all the mice during the probe trial day. G. Longevity of hTau and CgA-KO/hTau mice shown as a Kaplan-Meier plots (n= 20 per group). Values are average +/-SEM. p-values in A, B, D, E and F was calculated using Two-way ANOVA followed by Tukey’s multiple comparison test. p-values in G was calculated using Gehan-Breslow-Wilcoxon test. \**p* < 0.05, \*\**p* < 0.01, \*\*\**p* < 0.001, \*\*\*\**p* < 0.0001

KO/hTau mice demonstrated increased preference towards the new object, indicative of improved recognition memory compared to hTau mice (**Supplementary Fig 3C**). Additionally, the rotarod test revealed significantly improved motor function in CgA-KO/hTau mice across multiple trials, indicating rescued neuromuscular function (**Supplementary Fig 3D**). Collectively, these behavioral tests showcased the restoration of learning and memory-related activities upon CgA depletion in hTau, suggesting that CgA plays a pivotal role in the decline of cognitive function. Notably, the CgA-KO group outperformed the WT cohort in all behavioral tests, aligning with the hypothesis that CgA induces aging and that its removal rejuvenates mice beyond their biological age.

The median lifespan of CgA-KO/hTau mice extended to 13.22 months, compared to 10.12 months in hTau mice (**Fig 3G**), indicating a remarkable ∼30% increase in lifespan upon CgA depletion. Notably, 57% (n = 8) of CgA-KO/hTau mice exhibited relatively healthy lives until 12 months of age without discernible disease symptoms. Moreover, 23.6% (n = 5) of CgA-KO/hTau mice survived until 14.5 months, in contrast with the hTau mice, of which none had reach this age. Collectively, these findings underscore the robust impact of CgA depletion in the hTau mouse model in improving their cognitive function and mitigating premature mortality.

### CgA depletion partially protects glial activity in hTau mice

Next, we determined the potential role of CgA depletion in attenuating neuroinflammation, a key contributor to neurodegeneration and subsequent brain atrophy in hTau mice (*24*). We measured the levels of glial cell markers, specifically CD68 for microglia and GFAP for astrocytes, using IHC. Strikingly, the CA1 region of the hippocampi of CgA-KO/hTau mice had significantly lower levels of CD68 compared to their hTau littermates (**Fig 4A–B**). Simultaneously, cortical extracts from CgA-KO/hTau mice demonstrated diminished levels of proinflammatory cytokines, comparable to WT mice, in contrast to the elevated levels seen in hTau mice (**Fig 4C–F**; **Supplementary Fig 4a A–D**). We further found that the plasma proinflammatory cytokine levels were higher in hTau mice at both three and nine months of age (**Supplementary Fig 4E-K**) compared to CgA-KO/hTau mice. This suggests that removing CgA attenuates the systemic inflammatory response in hTau mice. Moreover, the anti-inflammatory cytokine IL-10 level was higher in CgA-KO/hTau than in hTau mice (**Supplementary Fig 4L**). On the other hand, GFAP levels showed a mild trend of decrease in CgA-KO/hTau mice compared to hTau mice (**Fig 4G – H**).

**Figure 4.**
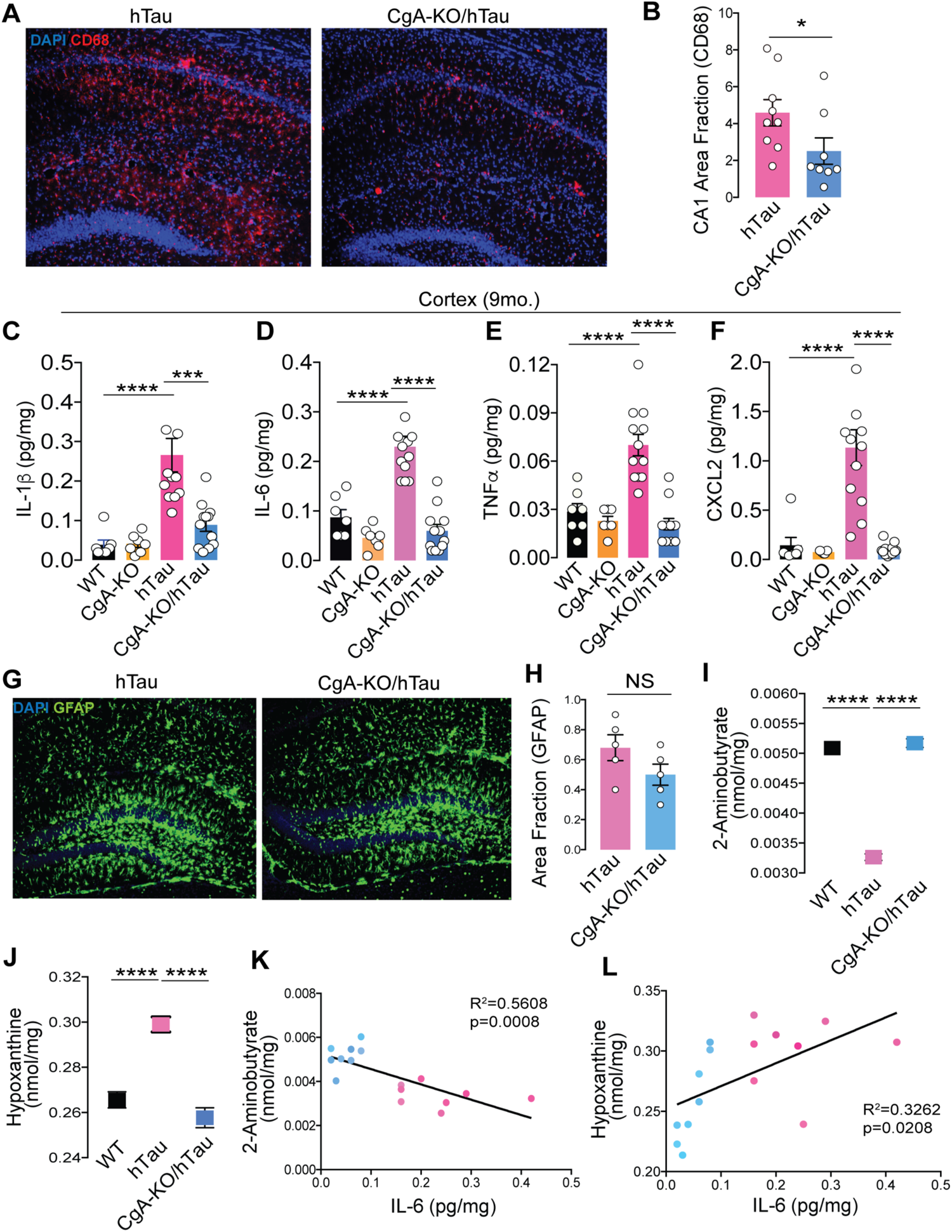
CgA-KO/hTau mice exhibit decreased microgliosis compared to hTau mice. A. Representative immunofluorescence (IF) images showing the levels of CD68 in the hippocampi (CA1 region) of hTau and CgA-KO/hTau mice. B. Quantification of the immunoreactive CD68 in CA1 as percentage of total CA1 area from IF images as shown in A. hTau (n=9), CgA-KO/hTau (n=8). C-F. Levels of cortical IL-1b (C), IL-6 (D), TNFa (E) and CXCL2 (F) in hTau (n=12), CgA-KO/hTau (n=12), WT (n-7) and CgA-KO (n=7). G. Representative IF images showing the levels of GFAP in the hippocampi (CA1 region) of hTau and CgA-KO/hTau mice. H. Quantification of GFAP fraction area calculated from IF mages as shown in G (n=5 per group). I-J. Levels of two metabolites, 2-aminobutyrate (I), and hypoxanthine (K) in the cortices of WT, hTau and CgA-KO/hTau (n= 8 per group). K-L. Pearson correlation analysis of 2-Aminobutyrate (K) and Hypoxanthine (L) with IL-6 level in the cortex of hTau (red dots) and CgA-KO/hTau (blue dots). Values are average +/-SEM. P-values in B was calculated using Mann-Whitney Test. p-values in C-F was calculated using one-way ANOVA followed by Tukey’s multiple comparison test. p-values in H was calculated using unpaired T-test. \**p* < 0.05, \*\**p* < 0.01, \*\*\**p* < 0.001, \*\*\*\**p* < 0.0001

Expanding our analysis to metabolic end products using gas chromatography followed by mass spectrometry, we identified minimal differences between WT, hTau, and CgA-KO/hTau mice for many metabolites, with noteworthy exceptions such as 2-aminobutyrate, hypoxanthine, and a few others (**Fig 4I–L**; **Supplementary Fig 4b A)**. Although these metabolites are relatively understudied in AD, existing literature suggests that 2-aminobutyrate may shield cells from oxidative damage by activating glutathione reductase (*25*) while hypoxanthine induces oxidative stress, mitochondrial abnormality, acetylcholinesterase activity, and neuroinflammation (*26–29*). We also observed changes in a few metabolites such as itaconate, tryptophan, sarcosine, spermidine, leucine, isoleucine, and methionine (**Supplementary Fig 4b B-H**) that either negatively or positively correlate with IL-6, matching their suggested roles in neuroinflammation and cognition (*30–32*). However, several other metabolites in CgA-KO/hTau remained similar to hTau mice. The complex interplay between CgA, neuroinflammation, and metabolic changes provides valuable insights into the multifaceted mechanisms underpinning tau-induced pathogenesis.

### CgA loss reverses transcriptomic changes in *hTau* mice

To elucidate the mechanism by which CgA depletion mitigates the pathogenic burden in hTau, we conducted bulk RNA sequencing on the cortices of three mouse genotypes (WT, hTau, and CgA-KO/hTau; n = 5 or 6 in each group; 3 males and 2-3 females). In comparing WT and hTau, 1,193 genes showed significantly higher expression and 869 were lower in WT than in hTau (log2(fold change) > 1.5, FDR < 0.01). Importantly, 417 of the upregulated genes (34.9%) and 293 of the downregulated genes (33.7%) in WT were similarly up- and downregulated in CgA-KO/hTau compared to hTau. These genes showing reversed expression in CgA-KO/hTau are represented in two clusters (**Fig 5A–D, F**). Gene ontology analyses of these clusters revealed significant enrichment in pathways involved in adenylate-cyclase-linked G-protein-coupled receptor (GPCR) signaling, ion transport, catecholamine transport, microtubule assembly, membrane potential, and exocytosis (**Fig 5E, G**). Given CgA’s role in catecholamine storage regulation and the influence of catecholamines on GPCR signaling (*11*), we determined the receptor genes in these pathways. The alpha-adrenergic receptor 1 (Adra1) family genes (α_1A_, α_1B_, α_1D_) were upregulated, while the Adra2 family genes (α_2A_, α_2B_, α_2C_) were downregulated in hTau compared to WT (**Fig 5A**; **Supplementary Fig 5A**), with no significant change in beta-adrenergic receptors (Adrb) across all the groups. RT-qPCR further confirmed altered gene expression of Adra1 and Adra2 in hTau, which were restored upon CgA deletion (**Supplementary Fig 5B–E**). Analysis of publicly available RNAseq data from control and AD patients demonstrated altered expression of Adra1a and Adra2a in Braak stages V–VI compared to Braak stages 0–I in parahippocampal gyri, indicating a correlation with disease progression (RNAseq data obtained from the Mount Sinai Brain Bank [MSBB] study; (**Supplementary Fig 5F–I**). No significant change in expression of Adra1b and Adra2b was observed in MSBB data sets, indicating species-specific effects. Functional analysis highlighted the interconnected role of Adra signaling in various cellular functions (catecholamine transport, neuropeptide signaling, second messenger signaling, and potassium-ion transport), underscoring its significance in tau pathology (**Supplementary Fig 5J**). Reports from other groups also described the importance of Adra1 in aggravating the disease pathology in the Aβ mouse model (*33*). Altogether, it is evident from our study that a significant alteration in the expression of Adra1 and Adra2 receptor classes and their downstream signaling cascades is strongly correlated with the pathogenesis.

**Figure 5.**
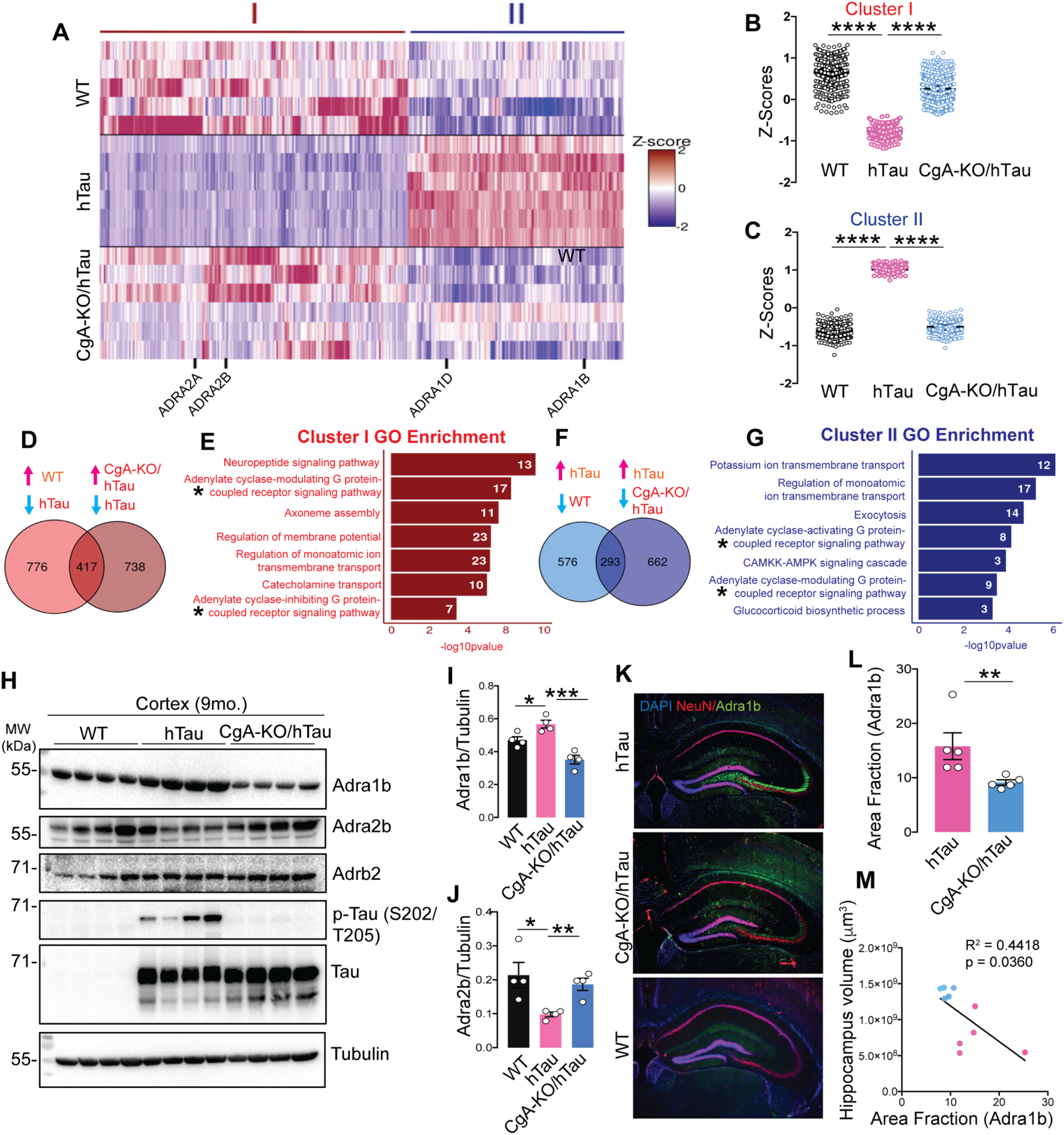
CgA deletion induces reprogramming of adrenergic receptor expression in tauopathy mice brain. A. Heatmap representation of normalized read Z-scores of shared upregulated or downregulated genes in WT (n=5) and CgA-KO/hTau (n=6) relative to hTau (n=6) (log2 fold-change > 0.5, p value < 0.01) from 9-month-old mice cortices. *Adra* gene family showing significant expression changes are highlighted. B. Z-score of hierarchical cluster I of WT, hTau and CgA-KO/hTau. C. Z-score of hierarchical cluster II of WT, hTau and CgA-KO/hTau. D. Venn diagram showing number of upregulated genes as compared in two groups (WT/hTau and CgA-KO/hTau). Expression of 417 genes overlaps in hTau vs. WT and hTau vs. CgA-KO/hTau. E. Gene ontology analysis for pathways enriched in shared genes in D. Number of genes involved in each pathway are shown in the bar. F. Venn diagram showing number of downregulated genes as compared in two groups (WT/hTau and CgA-KO/hTau). Expression of 293 genes overlaps in hTau vs. WT and hTau vs. CgA-KO/hTau. G. Gene ontology analysis for pathways enriched in shared genes in E. Number of genes involved in each pathway are shown in the bar. H. Representative WB showing Adra1b, Adra2b and Adrb2 protein levels in three mouse groups, WT, hTau and CgA-KO/hTau (n=4/group). Also shown are the levels of p-Tau (S202/T205), total hTau and tubulin (loading control). I. Quantitation showing relative levels of Adra1b in H. J. Quantitation showing relative levels of Adra2b in H. K. Representative IF images of Adra1b, NeuN and DAPI in the hippocampus slices of hTau, CgA-KO/hTau and WT mice. L. Image J Quantitation of IF images as in H showing relative levels of Adra1b in hTau and CgA-KO/hTau (n = 4/group). M. Pearson correlation between Adra1b area fraction and the hippocampus volumes of hTau (red dots) and CgA-KO/hTau (blue dots) mice. Values are average +/-SEM. p-values in B, C and I was calculated using one-way ANOVA (Sidak’s Multiple Comparison test). p-values in J and L was calculated using un-paired T-test. \**p* < 0.05, \*\**p* < 0.01, \*\*\**p* < 0.001, \*\*\*\**p* < 0.0001

Verification of protein levels for Adra1b and Adra2b in the cortex aligned with transcript levels. WB analysis demonstrated considerably lower Adra2b levels in the cortical extracts of hTau mice compared to WT and CgA-KO/hTau mice, while Adrb2 levels remained comparable (**Fig 5H–J**). IHC confirmed higher Adra1b protein levels in the CA3 cell body of hippocampal neurons in hTau mice compared to WT and CgA-KO/hTau (**Fig 5K–L**). Moreover, the increase in the area fraction of Adra1b is significantly correlated with the decrease in hippocampus volume (**Fig 5M**). Collectively, these findings suggest that CgA-driven exaggerated Adra1 and compromised Adra2 signaling constitute a key event in tauopathy and related dementia, including AD.

### Activated Adra1 signaling primes tau pathology in hTau mice

Adra1 receptors are intricately linked to G-protein signaling pathways, particularly through Gq, which triggers calcium flux. This calcium influx activates several kinases, including protein kinase C (PKC). Additionally, calcium can activate specific members of the adenylate cyclase family, leading to increased cAMP levels (*34*). In contrast, Adra2 receptors, serving as negative regulators in the adrenergic system, induce the inhibitory G protein Gi, blocking adenylyl cyclase and suppressing cAMP production (*35*). Aside from the Adra1/2 family members, CgA-KO/hTau reversed the altered expression of genes encoding phosphodiesterases (PDEs) in hTau compared to in WT mice. For instance, Pde1, Pde3, and Pde9 were expressed at lower levels in hTau mice than in WT and CgA-KO/hTau mice (**Supplementary Fig 6C**). The observed imbalance with elevated Adra1 and diminished Adra2 and PDE levels in hTau mice prompted an investigation into cAMP levels, a critical second messenger in the adrenergic system, within the cortex of mice. As anticipated, higher cAMP levels were observed in the cortices of hTau mice compared to those in WT and CgA-KO/hTau mice (**Supplementary Fig 6A–B**). Further analysis of patient samples revealed significantly higher cAMP levels in the cortices of Braak stage VI samples compared to Braak stage 0–II samples. These findings suggest that the upregulation of Adra1 signaling and downregulation of Adra2 signaling contribute to a signaling cascade maintaining elevated cAMP levels, thereby indicates probable activation of numerous tau kinases and fostering tau toxicity.

To explore whether dysregulated Adra1 signaling could induce tau pathology, tau phosphorylation and aggregation were assessed in OTSC treated with an Adra1 antagonist (Prazosin) or agonist (phenylephrine). OTSC transduced to express P301S human tau (AAV2-P301S-hTau) were exposed to K18 Tau fibrils in the presence or absence of Prazosin (**Fig 6A**). Three weeks post-treatment, WB analysis revealed significantly lower levels of pathological tau phosphorylation at residues S202, S396, and S404 in Prazosin-treated slices compared to untreated ones (**Fig 6B–D**). Additionally, IHC using the MC1 antibody showed reduced levels of tau aggregation in Prazosin-treated slices (**Fig 6E–F**). These findings strongly suggest that Adra1 signaling promotes the development of tau pathology. Conversely, in CgA-KO slices treated with an Adra1 agonist phenylephrine, levels of phosphorylated tau were increased (**Fig 6B–D**), and MC1+ tau aggregation was elevated compared to PBS-treated CgA-KO slices (**Fig 6E–F**). These data suggest that CgA-KO protects against tauopathy via inhibiting Adra1 signaling.

**Figure 6.**
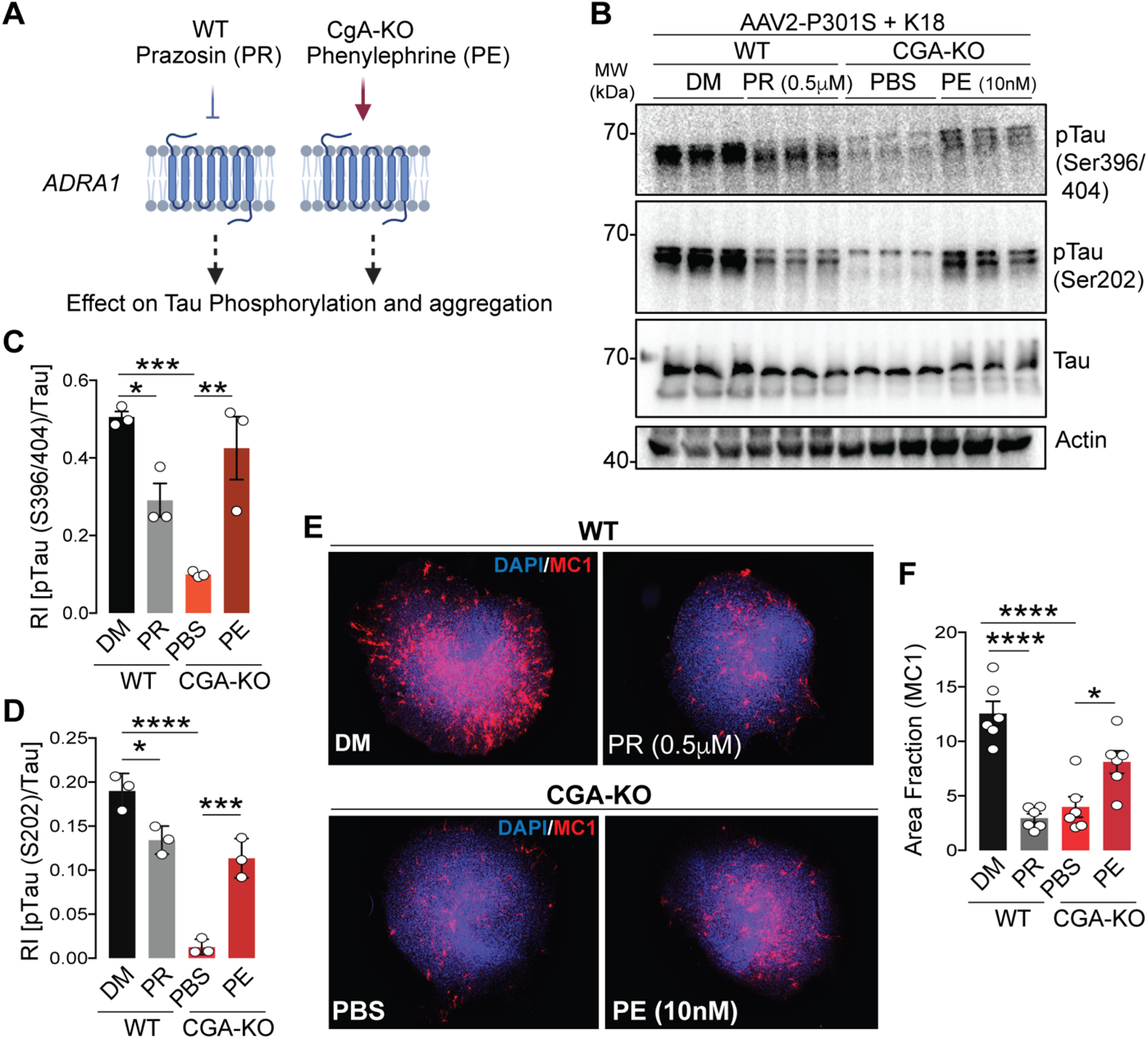
Activation and inactivation of alpha-1 adrenergic signaling regulates tauopathy. A. Schematics showing the rationale of Phenylephrine (Adra1 agonist) and Prazosin (Adra1 antagonist) treatment in WT and CgA-KO hippocampal OTSC to access the Tau aggregation ex-vivo. B. Representative WB showing pTau, total tau and actin (loading control) from OTSC lysates upon different treatment. C. WB quantification of Tau phosphorylation at S396/S404 relative to total tau levels (n=3). D. WB quantification of Tau phosphorylation at S202 relative to total tau levels (n=3). E. Representative IF images showing MC1+ Tau aggregates in WT (top) and CgA-KO (bottom) hippocampus OTSC treated with Prazosin (PR) and DMSO (DM) control (Top) and Phenylephrine (PE) and PBS control (Bottom), respectively. F. Image J quantification of MC1+ tau aggregates in OTSC as percentage of total slice area from IF images as shown in E. (n=6 per group) Values are average +/-SEM. p-values in C, E and F was calculated using One-way ANOVA. \**p* < 0.05, \*\**p* < 0.01, \*\*\**p* < 0.001, \*\*\*\**p* < 0.0001

### Epinephrine enhances tau phosphorylation and aggregation

The above results suggest that either EPI or NE or both are involved in Adra1-induced disease progression. Therefore, we assessed EPI and NE levels in the cerebrospinal fluid (CSF) and cortices of Braak stage I/II and stage VI patient samples. Although we found EPI levels were significantly higher in the CSF of Braak VI samples compared to Braak I samples, NE levels were found to be lower (**Fig 7A–C**; **Supplementary Fig 7A)**. Higher EPI levels were also found in the cortices and hippocampi of Braak VI patient samples compared to Braak I samples (**Fig 7C–D**). We also measured EPI and NE levels in the cortices of CBD patients’ samples and normal controls. Similar to AD samples, CBD samples showed higher EPI levels than in normal controls. NE levels did not differ between patients and controls samples. These results are consistent with prior reports indicating elevated EPI concentrations in the CSF of AD and dementia patients, correlating with disease progression, while the difference in NE levels between patients and nonpatients was not statistically significant (*36*). These findings suggest the potential involvement of CgA in the secretion and/or metabolism of EPI. Strikingly, both EPI and NE levels were elevated in the cortices of hTau mice compared to WT, CgA-KO, and CgA-KO/hTau mice, a trend similarly observed in the plasma (**Fig 7D**; **Supplementary Fig 7B**).

**Figure 7.**
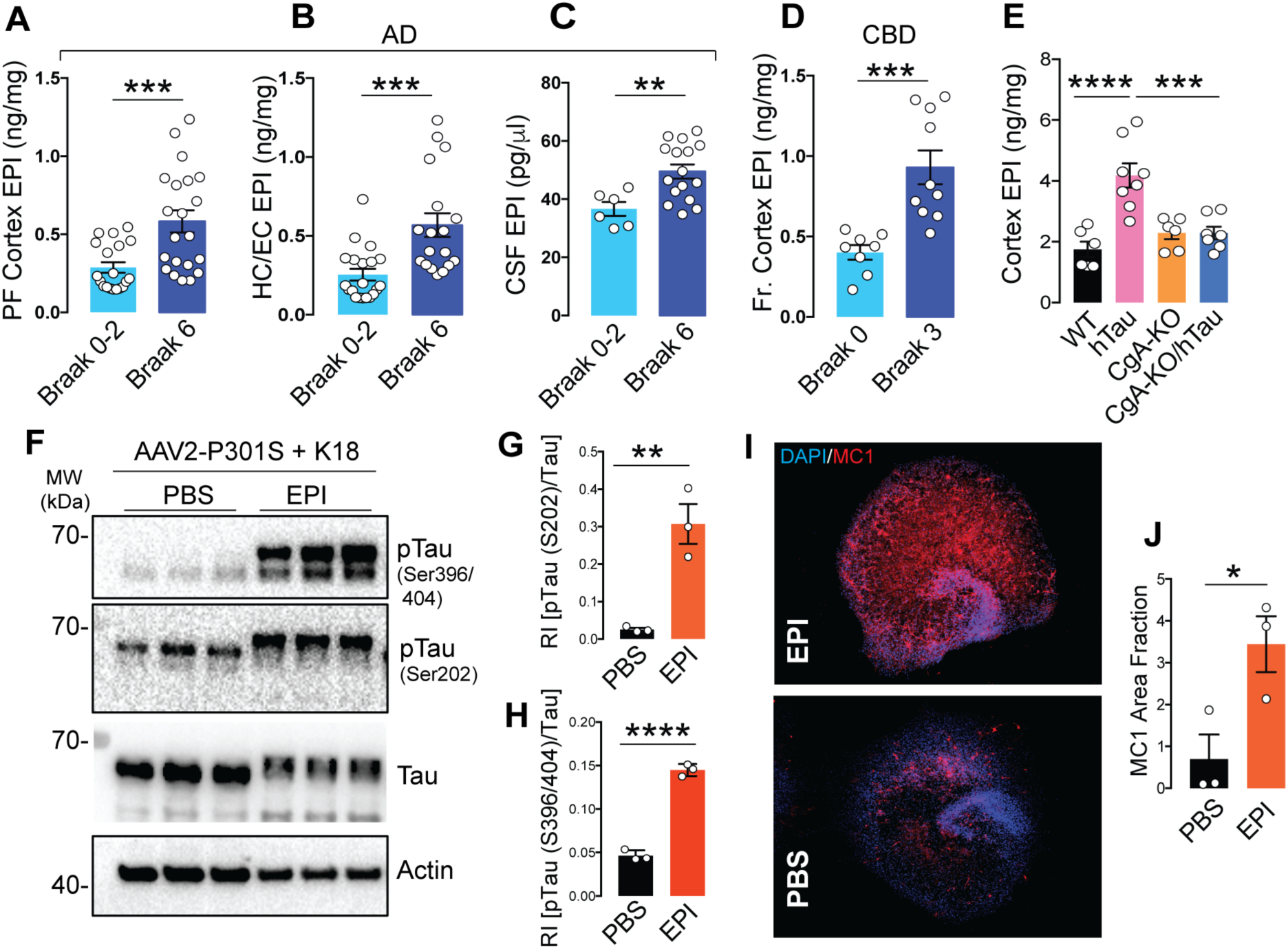
Elevated Epinephrine (EPI) levels in AD, CBD and hTau transgenic mice samples increases tauopathy in OTSC. A. Quantification of epinephrine (EPI) levels in the frontal cortex of AD patients from Braak stage 0-2 (early) and Braak stage 6 (late). B. Quantification of EPI levels in the hippocampus/Entorhinal cortex of AD patients from Braak stage 0-2 (early) and Braak stage 6 (late). C. Quantification of EPI levels in the CSF of AD patients from Braak stage 0-2 (early) and Braak stage 6 (late). D. Quantification of EPI levels in the pre-frontal cortex of CBD patients from Braak stage 0 (early) and Braak stage 3 (late) stages. E. Quantification of EPI levels in the cortex of 9-months old mice WT, hTau, CgA-KO, and CgA-KO/hTau (n=6-8 per group). F. WB showing pTau, total tau and actin (loading control) in hippocampus OTSC lysates upon EPI treatment. G. Quantification of WB of Tau phosphorylation at S202 relative to total tau (n=3). H. Quantification of WB of Tau phosphorylation at S396/S404 relative to total tau (n=3). I. Representative IF images showing MC1+ Tau aggregates in hippocampus OTSC treated with EPI (top) or PBS control (bottom). J. Image J quantification of MC1+ fraction in EPI and PBS treated hippocampus OTSC. Values are average +/-SEM. p-values in A, B, C, D, E, F and J was calculated using One-way ANOVA. \**p* < 0.05, \*\**p* < 0.01, \*\*\**p* < 0.001, \*\*\*\**p* < 0.0001

Given that neither EPI nor NE can cross the blood-brain barrier, these findings are particularly noteworthy. Since EPI levels exhibited enhancement in hTau mice and in CBD and AD samples, we focused specifically on EPI. To understand the impact of EPI on tau pathology, we explored whether EPI could induce tau phosphorylation and fibril formation in hippocampal OTSC. Remarkably, EPI significantly induced tau phosphorylation, indicating a pivotal role for aberrantly expressed brain EPI in the pathogenesis of tauopathy and AD (**Fig 7E-G**). IHC using the MC1 antibody confirmed that EPI also induced tau aggregate formation (**Fig 7H**). These observations underscore the potential contribution of dysregulated EPI levels to the development of tau pathology, further implicating a role for CgA in the intricate processes underlying in tauopathies including CBD and AD.

## Discussion

This study uncovers previously unexplored roles of CgA and EPI in the pathogenesis of tauopathies including AD, CBD and mouse models. We discovered that in the presence of CgA, mutant tau (TauP301S) transgenic mice have increased EPI levels and aberrant Adra signaling. Depletion of CgA normalized Adra signaling, reduced tau pathology and ameliorated tau-mediated neurodegeneration. This highlights the significance of aberrantly elevated EPI signaling as a key contributor tau pathogenesis in vivo. These findings align with the clinical observations in CBD and AD patients, where elevated EPI levels have been documented in the CSF (*36, 37*). Additionally, we observed increased EPI levels in other brain regions such as the cortex and hippocampus, along with an increase in Adra1 and a decrease in Adra2 transcripts in AD brains. Catecholamines such as EPI and NE, are co-stored with CgA and other molecules in brain storage vesicles (*38*). The release of these catecholamines from vesicles to nerve terminals and from terminals to synapses is controlled at different levels. Two key regulators are Adra2 and catestatin (CST), a peptide derived from CgA. Activation of Adra2 by EPI or NE in presynaptic neurons blocks their secretion through inhibition of calcium-mediated exocytosis of catecholamine storage vesicles (*39*). The CgA-derived peptide CST has been reported to inhibit catecholamine secretion from chromaffin cells (*40*). Therefore, it is likely that the lack of CgA processing into CST and Adra2 downregulation are related and are responsible for enhanced EPI secretion in the brain. This notion is supported by reports that EPI lowers the transcription of the Adra2a gene in rat astroglia by acting on Adra1 and Adrb and activating PKC and cAMP (*41, 42*). Although the reciprocal relationship of expression levels between Adra1 and Adra2 has not been shown in neurons, their presence in all brain cell types suggests EPI might exert its effect in neurons as well. Our observation that heightened Adra signaling is at least partly responsible for tau pathogenesis aligns with the previous finding that Adra1 antagonist alleviates memory loss in a mouse model (APP23) of AD (*43*). Future research will address the precise mechanism of EPI secretion in the brain.

Our observations parallel reports in the rhesus monkey model of AD linking increased Ca^2+^, cAMP, phosphorylated tau, and tangles (*44, 45*). However, excess NE, not EPI, was attributed to elevated Adra signaling (*35, 46*). Although excess NE could be pathogenic, our observations strongly suggest that the overactive EPI-Adra1 signaling pathway is key to tau phosphorylation and aggregation in brain regions like the hippocampus and cortex. Collectively, these observations implicate the dysregulation of Adra signaling, including upregulation of Adra1 and down regulation of Adra2, exacerbates tau pathogenesis .

The concentration of EPI in the brain represents only a fraction of the total EPI; the adrenal medulla is its primary source, acting as a hormone in peripheral organs. The intriguing question of how brain EPI functions as a neurotransmitter remains unanswered. The exclusive producers of EPI in the brain are the C1 neurons, predominantly located in the brainstem medulla, along with the C2/C3 neurons and the hypothalamus (*47*). These neurons express phenylethanolamine N-methyltransferase (PNMT), the enzyme responsible for EPI’s synthesis from NE (*48*). EPI is present in C1 nerve terminals that are projected to noradrenergic neurons in the locus coeruleus (LC) of the pons in the brainstem, functioning as an excitatory neurotransmitter (*49, 50*). LC is the sole source of noradrenergic neurons, playing a crucial neuroprotective role by facilitating communication with other brain regions. AD patients exhibit elevated PNMT levels in the cell body but decreased levels in the nerve ends of C1 neurons, indicating spaciotemporal deregulation of EPI synthesis in AD (*51, 52*). These published reports and our observations prompted us to propose that the surplus EPI produced in C1 neurons of TauP301S mice is released to postsynaptic noradrenergic neurons affecting homeostasis. Thus, C1 EPI might be the pathogenic trigger. Unregulated EPI can also impact glial cells, as they express Adra, potentially triggering inflammation. Similar EPI-induced inflammation has been observed in the lungs, where EPI activates cytokine expression in lung phagocytes (*53, 54*). Therefore, unchecked EPI may play a substantial role in microglia activation and neuroinflammation. Importantly, LC is the brain region affected by tau pathology at the earliest stage. The C1 nuclei, situated in the medulla oblongata beneath the pons in the brainstem, are responsive to various stressors such as inflammation, hypoglycemia, hypotension, and hypoxia, contributing to the maintenance of physiological homeostasis (*47, 55*). Given the associations between tauopathy/AD and inflammation, hyperglycemia, and hypertension and the pivotal role played by C1 neurons in regulating these stresses, it is plausible that EPI from C1 neurons plays a multifaceted role in both health and disease. Future research is necessary to unravel the intricate details of EPI’s role in the brain.

We further propose that CgA and the mutant Tau P301S protein form a direct association, initiating nonphysiological processes such as diminished axonal transport, EPI storage, synthesis, and release. This complex also hinders the processing of CgA into its bioactive peptides CST, which possesses anti-inflammatory properties, maintains insulin sensitivity, and inhibits catecholamine release (*40, 56, 57*). The absence of CST may contribute to the pathogenic process. Further studies are essential to elucidate the intricate relationship between CgA and Tau, determining how they combine to induce EPI levels. Subsequent experiments should also clarify the timing of EPI activation and the cell types where EPI is activated, shedding light on whether the EPI-Adra axis mediates the pathogenic process as an initiator or exacerbates an already-initiated process.

## Acknowledgments

This work was supported by grants AG072487 to GG, SKM, and XC; AG078635 to SKM and GG; AI163327 & GM085490 to GG; R01AG078185, R01AG074273 & UCSD Neurosciences start-up funds to XC. IL receives the following NIH grants: 5U01NS112010/807745, U01NS100610, R25NS098999; U19 AG063911-1 and 1R21NS114764-01A1. LT also receives supports from the Michael J Fox Foundation, Parkinson’s Foundation, Roche, AbbVie, Lundbeck, EIP-Pharma, Alterity, Novartis, and UCB. SJ was supported by an AFTD Holloway Postdoctoral Fellowship (2022-0002), and DM-M was supported by the AARF-D fellowship from the Alzheimer’s Association. The results published here are in whole or in part based on data obtained from the AD Knowledge Portal (adknowledgeportal.org). These data were generated from postmortem brain tissue collected through the Mount Sinai VA Medical Center Brain Bank and were provided by Dr. Eric Schadt from the Icahn School of Medicine at Mount Sinai. We thank Drs. Tapas Hazra and Tapan Biswas for sharing ideas, critiquing the work all through its course, and commenting on the manuscript. We also thank Drs. Malini Sen, Simpson Joseph, Balaji Krishnan, Tom Huxford and Sudhriti Ghosh Dastidar for commenting on the manuscript. We thank UCSD Shiley-Marcos Alzheimer’s Disease Research Center (ADRC) for providing AD postmortem brain tissues, blood and CSF samples, and Dr. Dennis Dickson for providing CBD postmortem brain tissue samples from the Jacksonville Mayo Clinic Brain Bank. This work includes data generated at the UC San Diego IGM Genomics Center utilizing an Illumina X Plus that was purchased with funding from a National Institutes of Health SIG grant (#S10 OD026929). We want to acknowledge **UCSD** School of Medicine **Microscopy Core** (Grant P30 NS047101).

## Contribution to work

SJ designed experiments, performed most experiments, analyzed data, and organized all figures. SS, DM-M, KT and YT performed experiments. GG designed experiments, organized figures, supervised the project, and wrote the manuscript with significant input from SJ and some help from SS. IL helped planning the CBD experiments, helped obtaining the patient samples and revised the manuscript. SKM designed experiments, performed experiments, analyzed some data, and supervised the project. XC designed experiments, helped in data analysis, supervised the project and revised the manuscript.

## Materials and Methods

### Animals

Chromogranin A knockout (CgA-KO) mice on a mixed background (50% 129/SvJ; 50% C57BL/6) were backcrossed to C57BL/6J mice for 10 generations to get CgA-KO mice in C57BL/6 background. CgA-KO mice were later inbred for 7 generations to establish a homogeneous mouse line. Those CgA-KO mice were backcrossed to B6C3F1/J mice for 4 generations to generate CgA-KO mice in B6C3F1/J background. These mice were crossed with hTau heterozygote mice (in B6C3F1/J background) to generate CgA-KO/hTau mice. Animals were kept in a 12-hour light/12-hour dark cycle. Mice have access to food and water ad libitum. Mice were fed a regular chow diet (NCD: 14% calorie from fat; LabDiet 5P00). All studies performed on animals were approved by the UCSD and Veteran Affairs San Diego Institutional Animal Care and Use Committee (IACUC) and conform to relevant National Institutes of Health guidelines.

### Genotyping

Mice were ear-tagged and tail-snips were collected. Genomic DNA from tail-snips were isolated using the Accustart Genotyping kit and amplified by PCR with Accustart Geltrack PCR supermix. Primers used for hTau genotyping are Forward (WT and mutant): 5’ TTG AAG TTG GGT TAT CAA TTT GG 3’, Reverse (WT): 5’ TTC TTG GAA CAC AAA CCA TTT C 3’, Reverse (Mutant): 5’ AAA TTC CTC AGC AAC TGT GGT 3’. Primers used for ChgA genotyping are Forward: 5’GTA GCA TGG CCA CTA CCC AG 3’ and Reverse: 5’ ATC CTT CAG AGC CCC TCC TT 3’.

### Hippocampal Slice Culture

Organotypic hippocampal slice cultures (OTSC) were performed with hippocampal slices obtained from postnatal day 7-8 pups of WT or CgA-KO mice as described previously (Croft & Noble, 2018). Pups were culled by decapitation, and the hippocampus was quickly dissected in oxygenated artificial cerebrospinal fluid (ACSF). Approximately 18-24 400 μm thick slices were cut using a tissue chopper, and these were cultured in Millicell culture inserts (Millipore) in 6 well plates (6-8 slices per insert). Culture media was changed every 2-3 days and the slices were harvested after 21 or 28 days. The composition of ACSF was 125mM NaCl, 2.4mM KCl, 1.2mM NaH_2_PO_4_, 1mM CaCl_2_, 2mM MgCl_2_, 25mM NaHCO_3_, and 25mM Glucose. The composition of OTSC culture media was 25mM HEPES, 133mM NaCl, 31mM NaHCO_3_, 45mM D-Glucose, 5mM KCl, 3.3mM MgSO_4_, 4.5mM CaCl_2_, 1mM Na_2_HPO_4_, 0.5mM Ascorbic Acid, 1,25ml Pen-Strep, 2.5ml Glut-Max, 62.5ml Heat Inactivated Horse Serum and 82.5μl Insulin.

### In vitro fibril-induced tau spreading assay

Hippocampal slices were transduced with AAV viral vector expressing P301S human tau (AAV2-smCBA-human_P301S_Tau-WPRE; 2x10^10^/mL) for 24h. AAV was removed from culture media and K18 (preformed tau fibrils; 1.5 μg/mL) were added before incubation for 72h. Subsequently, fresh media without or with Phenylephrine:10nM, Prazosin:0.5μM, or Epinephrine: 10nM, was applied as indicated. Slices were harvested on day 21 for Western blot analysis and immunohistochemistry.

### In vivo fibril-induced tau spreading assay

Mice were anesthetized with inhalation of 2% isoflurane for the duration of surgery and secured on a stereotaxic frame (Kopf Instruments). 3-month-old hTau and CgA-KO/hTau mice were injected stereotaxically at a rate of 0.5μl/min, with 2μL of 5 mg/ml K18 PFF into the CA1 region of the left hippocampus. The coordinates for injection were anterior-posterior − 2.5, medial-lateral + 2.0, dorsal-ventral −1.8. Mice were sacrificed 6 weeks after injection and transcardially perfused with PBS for immunohistochemistry analysis of tau aggregation (MC1+) in the ipsilateral and contralateral sides.

### K18 purification and *invitro* Fibril formation

The repeat domains (K18) of P301L mutant Tau with a Myc Tag was expressed in *E.coli* BL21 (DE3) strain. NaCl (500mM) and betaine (10mM), a small molecule chaperone, were added before induction with IPTG (200 μM) at 30°C for 3.5h. Cells were resuspended in lysis buffer (20mM MES, pH 6.8, 1mM EGTA, 0.2mM MgCl_2_, 1mM PMSF, 5mM DTT), and passed through a microfluidizer for lysis. After lysis, 5M NaCl was added to a final concentration of 500mM followed by heating at 90°C for 20min. was clarified by centrifugation and was dialyzed overnight (Buffer A: 50mM NaCl, 1mM MgCl_2_, 0.1mM PMSF, 2mM DTT, 1mM EGTA, 20mM MES pH 6,8) to obtain the Tau-containing supernatant. Subsequently, cation exchange (HiTrap SP HP, 5ml column from Cytiva) was performed to purify K18. The degradation products and other impurities were removed by running 15% elution buffer (50mM NaCl, 1mM MgCl_2_, 0.1mM PMSF, 2mM DTT, 1mM EGTA, 20mM MES pH 6,8, 1M NaCl).

Subsequently, 15-60% gradient (NaCl gradient from 50mM to 1M) of NaCl was used to obtain pure K18, which was concentrated and further dialyzed against assay buffer (PBS pH 7.4, 1mM DTT and 2mM MgCl_2_). A 10μM concentration of Tau was incubated in assay buffer for 36h at 37°C) upon adding 44μg/ml Heparin. The aggregates formed were snap frozen at -80°C till further use.

### Transmission Electron Microscopy

Deeply anesthetized mice were flushed with a pre-warmed (37°C) Hank’s balanced salt solution (HBSS), with calcium and magnesium by perfusion with freshly prepared pre-warmed (37°C) fixative containing 2.5% glutaraldehyde and 2% paraformaldehyde in 0.15 M cacodylate buffer using a peristaltic pump. The hippocampus and cortex were dissected and postfixed in 1% OsO4 in 0.1 M cacodylate buffer. The tissues were stained en bloc with 2-3% uranyl acetate and dehydrated in graded ethanol series. Sections of 50-60 nm thickness were cut, stained with 2% uranyl acetate and Sato’s lead stain. Grids were imaged with a JEOL JEM1400-plus TEM (JEOL, Peabody, MA) attached to a Gatan OneView digital camera with 4k x 4k resolution (Gatan, Pleasanton, CA).

### Measurement of catecholamines

Cortical and plasma catecholamines were measured upon separation by an Atlantis dC18 column (100A, 3 µm, 3 mm x 100 mm) on a ACQUITY UPLC H-Class System attached to an electrochemical detector (ECD model 2465; Waters Corp, Milford, MA) as described previously. The mobile phase (isocratic: 0.3 ml/min) contained 95:5 (vol/vol) of phosphate-citrate buffer and acetonitrile. An internal standard 3.4-dihydroxybenzylamine (DHBA 400 ng) was added to the cortical homogenate in 0.1N HCl. The ECD was set at 500 pA for determination of brain catecholamines. For determination of CSF catecholamines, DHBA (2 ng) was added to 150 µl CSF and adsorbed with ∼15 mg of activated aluminum oxide for 10 min in a rotating shaker. After washing with 1 ml water, adsorbed catecholamines were eluted with 100 µl of 0.1N HCl. The ECD was set at 500 pA for determination of CSF catecholamines. The data were analyzed using Empower software (Waters Corp, Milford, MA). Catecholamine levels were assessed using the internal standard. Catecholamine levels were provided in nM (CSF) or ng/mg protein. For determination of plasma catecholamines, DHBA (2 ng) was added to 150 μl plasma and adsorbed with ∼15 mg of activated aluminum oxide for 10 min in a rotating shaker. After washing with 1 ml water adsorbed catecholamines were eluted with 100 μl of 0.1N HCl. The ECD was set at 500 pA for determination of plasma catecholamines. The data were analyzed using Empower software (Waters Corp, Milford, MA). Catecholamine levels were normalized with the recovery of the internal standard. Plasma catecholamines were expressed as nM.

### Measurement of cytokines

Cortices from hTau, CgA-KO/hTau, WT and CgA-KO were homogenized in PBS followed by centrifugation at 12500 RPM for 30 min at 4°C. The supernatant was collected and used in ELISA. Cytokines were measured using U-PLEX mouse cytokine assay kit (Meso Scale Diagnostics, Rockville, MD) following the manufacturer’s protocol in a MESO SECTOR S 600MM Ultra-Sensitive Plate Imager. The levels were presented in pg/mg protein. Plasma (25 µl) cytokines were measured using U-PLEX mouse mouse cytokine assay kit (Meso Scale Diagnostics, Rockville, MD).

### Behavioral Studies

Behavioral studies including, Morris Water Maze, Novel Object Recognition and Rotarod test for all four cohorts of mice were performed in UCSD behavior core facility.

*Morris Water Maze (MWM) test*: MWM test is a long-term memory test where a chamber is filled with opaque water and there is a submerged platform under the surface. Training was carried out in the period of day1 to day7 for the hidden platform where mice were guided to reach the platform and trial was ended if they reached and are stationary on the platform for 5 seconds. Both distance and time were recorded for each mouse. On day8, the mice were dropped 180° opposite of the location of the platform but without the platform in the chamber. The number of times each mice entering the correct zone was recorded. Separately, the hidden platform was provided, and the time taken by each mouse to reach the platform was recorded and represented as latency.

*Novel Object Recognition (NOR) test*: In NOR, the mice was left to spend with time with a known object for 180 seconds on day1. The time for which the mouse explored the unknown object was recorded. On day2, a new object is introduced in addition to the old object. It is expected that the mouse will spend more time with the new introduced object for curiosity, if it can remember the old object. The total time spend by mice with the new object on day2 determines the degree of memory loss and represented as (Time with new object X 100)/Time with new + old object.

*Rotarod test*: Rotarod test was performed to evaluate the motor functions in mice. In this test there was a horizontal rod which rotates around its axis and the mice must co-ordinate with the movement so that they don’t fall off. The time for which they can stay on the rod determines their motor function. On day 1 each mice undergo five trials on the rotarod. On day 2, each mice undergo seven trials and time from each trial is plotted and compared.

### Tau Seeding Assay

Human Tau RD P301S FRET biosensor expressing HEK 293 cells (ATCC CRL-3275) were plated on Millipore EZ chambered slide at a confluency of 1 X 10^3^ Cells / well in DMEM Complete media (DMEM with 10%FBS+100 μg penicillin and streptomycin). After 16h, the media is replaced with OptiMEM and cells were treated with 2 μg lysate from CgA-KO/hTau and hTau cortex and 4μl Lipofectamine-3000. Post 24h of treatment the media is replaced with complete DMEM. After 24h the media was removed and cells were fixed with 4% Paraformaldehyde and stained with DAPI. Cells were mounted and images were taken using Keyence Fluorescence Microscope under 40X lens. Four biological replicates per group were taken and 4 images each group were analyzed by ImageJ for quantification.

### Immunohistochemistry

Isofluorane was used to anesthetize the mice followed by trans-cardial perfusion with PBS. The brain was dissected out and kept in Zn-Formalin for 48hr. After that, Zn-Formalin is replaced with 30% Sucrose incubated for 72hr at 4°C. The brains underwent coronal sectioning of 30μm thick by sliding freezing microtome (Epredia). The sections were kept at -20°C in cryoprotectant. Approximately 7-8 sections were taken for each animal covering from anterior to posterior region of hippocampus. The sections were washed thoroughly (6X 10 min wash) with 1X PBS and then incubated with primary antibody in PBS containing 0.4% Triton-X for 24hr at 4°C. Following which the primary antibody is removed and washed with 1X PBS for 3X 15min. Subsequently, the secondary antibody and DAPI (1:2000) were added and incubated for 1hr at room temperature followed by 3X 15min PBS wash. Next, the sections were mounted on glass slides with Fluoromount G. The stained slides were imaged under Keyence Fluorescence Microscope. Primary antibodies used: AT8 (1:500), MC1 (1:500), CD68 (1:700), GFAP (1:500), ADRA1B (1:500), CgA (1:400). Secondary antibodies used: Anti-mouse Alexa Fluor 568 (1:300),

Anti-rabbit Alexa Fluor 488 (1:300), Strep-Alexa Fluor 568 (1:500).

Paraffin-embedded brain slices pre-mounted on slides from ADRC were de-paraffinized in xylene for 10 mins, followed by rehydration with sequential incubation in 100%, 90%, 70% EtOH, 5 mins each. Sections were placed in Citrate buffer (pH6) solution in a pressure cooker at low pressure for 10 minutes for antigen retrieval. The slides were blocked (2.5% Normal Goat Serum for 1hr at room temperature) and incubated with primary antibody for overnight at 4°C, followed by 3X PBS wash for 15 min each. Next, a secondary antibody and DAPI (1:2000) were added and incubated for an hour at room temperature. After this, slides were dried and mounted with Fluoromount G. Images were taken with the Keyence Fluorescence microscope.

### Nissl Staining, Volumetric analysis and DG and CA1 thickness measurement

Investigators were blinded to the genotypes or treatment of the mice. For quantification of hippocampal volume, mice hemibrains were cut at 30 μm coronally, and all hippocampi, including sections, were collected. Brain sections were mounted on microscope slides (Fisher Scientific) in an anterior-to-posterior order, starting from the section where the hippocampal structure first becomes visible (first section) to the section where hippocampal structure just disappears (last section). Mice with missing sections were excluded from the analyses, a pre-established criterion. Mounted brain sections were dried at room temperature for 24 h and stained with cresyl violet (Nissl staining). After rehydrating with a quick wash in distilled water, sections were stained in FD Cresyl Violet Solution for 3 min. Next, sections were dehydrated in increasing ethanol concentrations and differentiated in 95% ethanol with 0.1% glacial acetic acid. After the final dehydration in 100% ethanol, sections were cleared in xylene and mounted with DePeX mounting media (VWR). Images were acquired with a Keyence BZ-9000 microscope. Hippocampal volume was estimated using ImageJ (NIH) Volumest plugin (http://lepo.it.da.ut.ee/∼markkom/volumest/). To measure the thickness of CA1 pyramidal cell layer and dentate gyrus granule cells layer, a straight line perpendicular to the length of cell layers at fixed locations was drawn and measured using ImageJ.10–12 hippocampal-containing sections were typically used for each analysis.

### Western Blotting

15mg Tissue was incubated in RIPA buffer (150mM NaCl, 1% Triton-X, 12mM Na-deoxycholate, 0.1% SDS, 50mM Tris pH 8.0, 25mM EDTA, 1mM PMSF, 5% Glycerol, 50mM NaF, 1mM Na3VO4, 1mM Na_4_P_2_O_7_, 25mM β-Glycerolphosphate, 50mM DTT and PIC) for 10 min. Homogenization was done by handheld motor homogenizer, followed by centrifugation at 12500 rpm for 30min at 4°C. The supernatant was collected and protein concentration was measured using Bradford Protein Assay Reagent. 15μg protein was loaded in each lane of 12% Tris-Glycine SDS Gel and post-running protein was transferred to PVDF membrane followed by blocking with 5% BSA for 2hrs. Primary antibody incubation was done at 4°C. Antibodies and dilution used as follows: 1°CgA (1:4000), 1°AT8 (1:3000), 1°PHF1 (1:4000), 1°CP13 (1:2000), 1°HT7 (1:8000), 1°Actin (1:8000), 1°ADRA1B (1:2000), 1°ADRA2B (1:2000), 1°ADRB2 (1:2000), 1α-Tubulin (1:6000), 2°Anti-Mouse-HRP (1:10000), 2°Anti-Rabbit-HRP (1:10000), Strep-HRP (1:6000).

### cAMP measurement

The cortex tissue samples from mice or human were lysed in PBS along with PDE inhibitor (IBMX) and PIC (SIGMA) with homogenizer followed by centrifugation at 12500 rpm for 20min at 4°C. The cAMP level in the supernatant was analyzed using Perkin Elmer cAMP measurement kit as described in the manual. Briefly, the tissue lysate was incubated with anti-cAMP acceptor beads in corning dark 384 well plate and incubated for 30min at room temperature in dark. Subsequently, biotinylated cAMP/streptavidin beads were added, and incubated at room temperature in dark for 1h. Excitation was performed at 680nm wave-length light and emission was measured at 550nM to determine the cAMP concentration based on the standard curve.

### Metabolite Analysis

Frozen brain samples (20-40 mg) were transferred to 2-mL tubes containing 2.8 mm ceramic beads (Omni International) and 0.45 ml ice-cold 50% methanol/ 20 µM L-norvaline was added.

Tubes were shaken (setting 5.5) for 30 s on a Bead Ruptor 12 (Omni International), quickly placed on ice, and frozen at -80°C overnight. Thawed samples were centrifuged at 15,000 x g for 10 minutes at 4°C. The supernatant was then transferred to a new tube, mixed with 0.225 ml chloroform, and centrifuged at 10,000 x g for 10 minutes at 4°C. This produced a two-phase separation. Portions (200 µl) of the top phase, along with 7 dilutions of a mixture of standards (see attached SRM sheet for list of standards), were dried (Speedvac) for analysis of polar metabolites. Dried samples and standards were derivatized with 30 µl isobutylhydroxylamine (TCI, 20 mg/ml in pyridine, 80°C for 20 min) followed by addition of 30 µl MTBSTFA (Soltec) and further incubation at 80°C for 60 min, before transfer to vials for GC-MS analysis. Samples and standards were analyzed by GC-MS using a TG-SQC column (15 m x 0.25 mm x 0.25 µm, Thermo) installed in a Thermo Scientific TSQ 9610 GC-MS/MS. The GC was programmed with an injection temperature of 250°C and a 0.8 µl injection with 1/5 split. The GC oven temperature was initially 130°C for 4 min, rising to 250°C at 6°C/min, and to 280°C at 60°C/min with a hold at the final temperature for 2 min. GC flow rate with helium carrier gas was 50 cm/s. The GC-MS interface temperature was 300°C and (electron impact) ion source temperature was 200°C, with 70 eV ionization voltage. Standards were run in parallel with samples. Metabolites in samples and standards were detected by MS/MS using precursor and product ion masses, and collision energies shown in the attached table (argon was used as the collision gas). Sample metabolites were quantified with calibration curves based on the standards (in Thermo Chromeleon software) and further data processing to adjust for the recovery of the internal standard (norvaline) and for the relative quantities of metabolites in the standards was done in MS Excel.

### RNA extraction and qPCR

RNA was isolated using TRIzol reagent and extracted further with phenol-chloroform method. RNA concentration was measured using Thermo Nanodrop and cDNA was synthesized from 500ng of total RNA using Maxima RT cDNA synthesis kit. SYBR green containing NEB Luna qPCR mix was used to set up the qPCR in Bio-Rad qPCR machine.

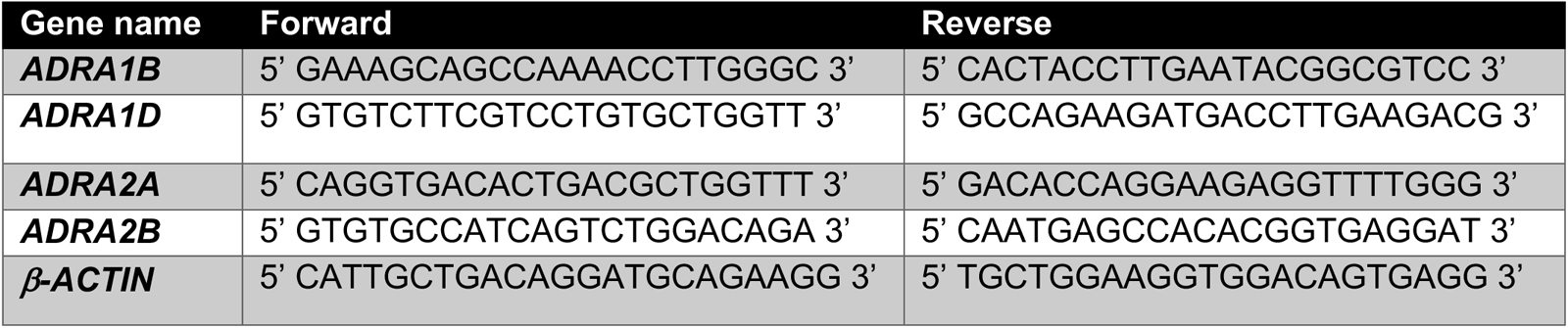

### RNA-seq library preparation and analysis

Total RNA was isolated from prefrontal cortex of 8-month-old mouse using RNeasy kit (Qiagen). RNA amount was quantified by ‘Nanodrop’ spectrophotometer and integrity was assessed by TapeStation (Agilent). Complementary DNA libraries were prepared from 500 ng of total RNA using the mRNA HyperPrep Kit (KAPA) according to manufacturer’s recommendations with Unique Dual-Indexed adapters (KAPA). Libraries were PCR-amplified for 10 cycles and quality was assessed by Tapestation. Libraries were then quantified by Qubit 2.0 fluorometer (Thermo), pooled, and analyzed by paired-end 100 sequencing on the NovaSeq 6000 platform (Illumina) at UCSD IGM Core.

### RNA-Seq Analysis

RNA samples were first assessed for Sequencing quality by FASTQC and libraries with high quality reads were trimmed using Trimmomatic (v0.39). Reads were mapped to the mm10 mouse reference genome using STAR (v2.7.10b) and transcripts were quantified by StringTie (v1.3.6). Transcript counts were imported to R using prepDE.py3 for normalization and differential expression was analyzed by DESeq2 (v1.42.0). Gene clusters among differentially expressed genes were identified as common genes in comparisons of WT vs hTau and CgA-KO/hTau vs hTau with log2 fold-change > +/-0.5 and p value < 0.01. For gene ontology analysis, the enrichGO function from clusterProfiler (v4.10.0) with the org.Mm.eg.db (v3.18.0) was used with BH parameter for p value adjustment.

### Human Transcriptomics Data Analysis

For human transcript analysis, normalized data of gene expression in parahippocampal gyrus (Brodmann area 36) was obtained from Mount Sinai Brain Bank (MSBB) through the Synapse platform (syn16795937).

| Study | Brain Region | Cohort | BRAAK Stage |  |  |  |  |  |  |  | # Control<br>(BRAAK 0-1) | # AD<br>(BRAAK 5-6) | # filtered genes |
| --- | --- | --- | --- | --- | --- | --- | --- | --- | --- | --- | --- | --- | --- |
|  |  |  | 0 | 1 | 2 | 3 | 4 | 5 | 6 | Total |  |  |  |
| Mount Sinai Brain Bank | Parahippocampal gyrus (BM36) | MSBB_BM36 | 10 | 24 | 35 | 40 | 23 | 24 | 65 | 221 | 34 | 89 | 23201 |

