## Supplemental Figures for "Chromogranin A (CgA) Deficiency Attenuates Tauopathy by Altering Epinephrine–Alpha-Adrenergic Receptor Signaling"

**Supplementary Figures:**

**Supplementary Fig 1**
**CgA co-localize with aggregated Tau (MC1) and CgA depletion in OTSC reduces Tau seeding**

A-B. Pearson correlation of pTau/Tau and CgA/Actin in AD (A) and CBD (B) samples.
C-D. Representative image of IHC of CgA in WT and hTau mice hippocampus (C) with quantification (D).
E-F. Co-localization of CgA (red) and MC1-detected aggregated Tau (green) in three different Braak 6 (E) and one Braak 1 (F) hippocampi.
G. Schematic of organotypic slice culture (OTSC) generation, treatment and imaging.
H. WB showing reduced pTau in AAV2-hTau (P301S) transduced CgA-KO slices compared to WT.
I-J. Densitometric quantification of phospho-Tau (Ser396/404, Ser202) between K18-treated and AAV2-hTau (P301S) transduced CgA-KO and WT slices.
K. Representative IF images using MC1 antibody showing tau aggregates in organotypic hippocampal slices from WT (top) and CgA-KO (bottom) mice transduced with AAV tau P301S and treated with K18 fibrils.
L. Image J quantification of tau aggregates as a ratio of area fractions in ipsilateral versus contralateral from images as represented left (n = 6).
p-values in D was calculated using unpaired T-test. P-values in K-L was calculated using one-way ANOVA (Sidak's Multiple Comparison Test).

**Supplementary Fig 2**
**Rescue of tau pathology in CgA-KO/hTau mice.**

A-B. Image J quantification of the thickness of DG (A) and CA1(B) in WT (n=6), hTau (n=14) and CgA-KO/hTau mice (n=11).
C-E. Densitometric quantification of WB of pTau (C & D) and PSD95 (E) in CgA-KO/hTau and hTau mice by image J (n=6).
F-G. Representative IHC images of p-Tau (Ser 202/Thr205) with quantification in CgA-KO/hTau and hTau mice (n=5).
H-I. Quantification of MC1+ Tau aggregates in ipsilateral (H) and contralateral (I) side of the brain.
J-K. Transmission electron microscopy (TEM) image showing synapses in CgA-KO/hTau and hTau mice (J), quantified in (K) as a function of area measured in $\mu m^2$.
L-M. TEM images showing synaptic vesicles in CgA-KO/hTau and hTau depicted by TEM images and quantification (M).

**Supplementary Fig 3**
**Improved cognitive and motor function in CgA-KO/hTau mice compared to hTau mice.**

A. Time taken to reach the platform in each day of MWM by all four mice group [WT (n=18), CgA-KO (n=16), hTau (n=19) and CgA-KO/hTau (n=20)].
B. Body weight for all four mice groups (WT, CgA-KO, hTau and CgA-KO/hTau). [WT (n=19), CgA-KO (n=20), hTau (n=16) and CgA-KO/hTau (n=20).
C. Novel Object Recognition scores of CgA-KO/hTau and hTau mice. [WT (n=15), CgA-KO (n=16), hTau (n=19) and CgA-KO/hTau (n=20)].
D. Time before falling in the Rotarod test for all four mice groups [WT (n=18), CgA-KO (n=16), hTau (n=20) and CgA-KO/hTau (n=20)].

P-values in A, B, C and D were calculated using Two-way ANOVA followed by Tukey's multiple comparison test. *$p < 0.05$, **$p < 0.01$, ***$p < 0.001$, ****$p < 0.0001$

**Supplementary Fig 4a**
**Neuroinflammation and systemic inflammation were reduced in CgA-KO/hTau mice compared to hTau mice.**
A-D. Inflammatory cytokine levels in the cortex of WT (n=7), hTau (n=12), CgA-KO (n=7) and CgA-KO/hTau (n=12) mice at 9 months of age (mo).
E-L. Plasma inflammatory cytokine levels at 3 mo and 9 mo age in WT, hTau, CgA-KO and CgA-KO/hTau mice. [3 mo; WT (n=6), CgA-KO (n=5), hTau (n=6), CgA-KO/hTau (n=5) and 9 mo; WT (n=5), CgA-KO (n=5), hTau (n=7), CgA-KO/hTau (n=7)]
P-values in C-F was calculated using one-way ANOVA followed by Tukey's multiple comparison test. *$p < 0.05$, **$p < 0.01$, ***$p < 0.001$, ****$p < 0.0001$

**Supplementary Fig 4b**
**Alterations of levels of metabolites in CgA-KO/hTau mice compared to hTau mice and correlation with inflammatory cytokine IL6.**
A. Difference in polar metabolites in CgA-KO/hTau, hTau and WT pups. N=8.
B-H. Pearson Correlation between polar metabolites and inflammatory cytokine (IL6) in cortex.

**Supplementary Fig 5**
**Validation of transcriptional changes of genes identified in RNA-seq of WT, hTau and CgA-KO/hTau hippocampus.**
A. Normalized counts of class I and class II alpha-adrenergic receptors in WT, hTau and CgA-KO/hTau hippocampus.
B-E. qPCR of genes in WT, hTau and CgA-KO/hTau.
F-I. Analysis of alpha-1 adrenergic receptor transcript level in Braak stage 1 and Braak stage 6 patients (parahippocampal gyrus) from AMP-AD database.
J. Network analysis of the pathway involved depicting adrenergic signaling pathway as an important one linked with catecholamine transport, neuropeptide signaling and potassium ion transmembrane transport.
p-values in A, E, F and H were calculated using unpaired T-test. p-values in B-D was calculated using one-way ANOVA (Sidak's multiple comparison test). *$p < 0.05$, **$p < 0.01$, ***$p < 0.001$, ****$p < 0.0001$.

**Supplementary Fig 6**
**Enhanced cAMP and Phosphodiesterase (PDE) level in CgA-KO/htau mice**.
A. Decreased cAMP level in the cortex of CgA-KO/hTau mice compared to hTau mice. n=10.
B. Increased cAMP level in Braak stage 6 patient (n=13) cortex lysates compared to Braak stage 1-2 (n=13).
C. RNA-seq analysis of differentially expressed genes (DEGs) showing the expression of different phosphodiesterases (PDEs) across the three genotypes (WT, hTau and CgA-KO/hTau).
p-values in A was calculated using Brown-Forsythe Test. p-values in B was calculated using unpaired T-test. *$p < 0.05$, **$p < 0.01$, ***$p < 0.001$, ****$p < 0.0001$

**Supplementary Fig 7**

**Catecholamine levels in AD and CBD patient cortex and in CgA-KO/hTau mice plasma.**

A-C. Nor-Epinephrine (Nor-EPI) levels in Braak 6 and Braak 0-2 AD prefrontal cortex (A), hippocampus/entorhinal cortex (B) and CSF (C). cortex [Braak 0-2 (n=19), Braak 6 (n=21)], hippocampus [Braak 0-2 (n=18), Braak 6 (n=20)]

D. Nor-EPI levels in CBD pre-frontal cortex. Braak 0 (n=8), Braak 3 (n=10).

E. Nor-EPI levels in cortex of WT (n=6), hTau (n=7), CgA-KO (n=6) and CgA-KO/hTau (n=7) mice.

F. Nor-EPI levels in plasma of WT (n=26), hTau (n=32), CgA-KO (n=26) and CgA-KO/hTau mice (n=32).

G. EPI level in plasma of WT (n=26), hTau (n=32), CgA-KO (n=26) and CgA-KO/hTau mice (n=32).

p-values in A-D was calculated using unpaired T-test. p-values in E, F and G was calculated using Two-way ANOVA followed by Tukey's multiple comparison test. *$p < 0.05$, **$p < 0.01$, ***$p < 0.001$, ****$p < 0.0001$.

### Supplementary Fig 1

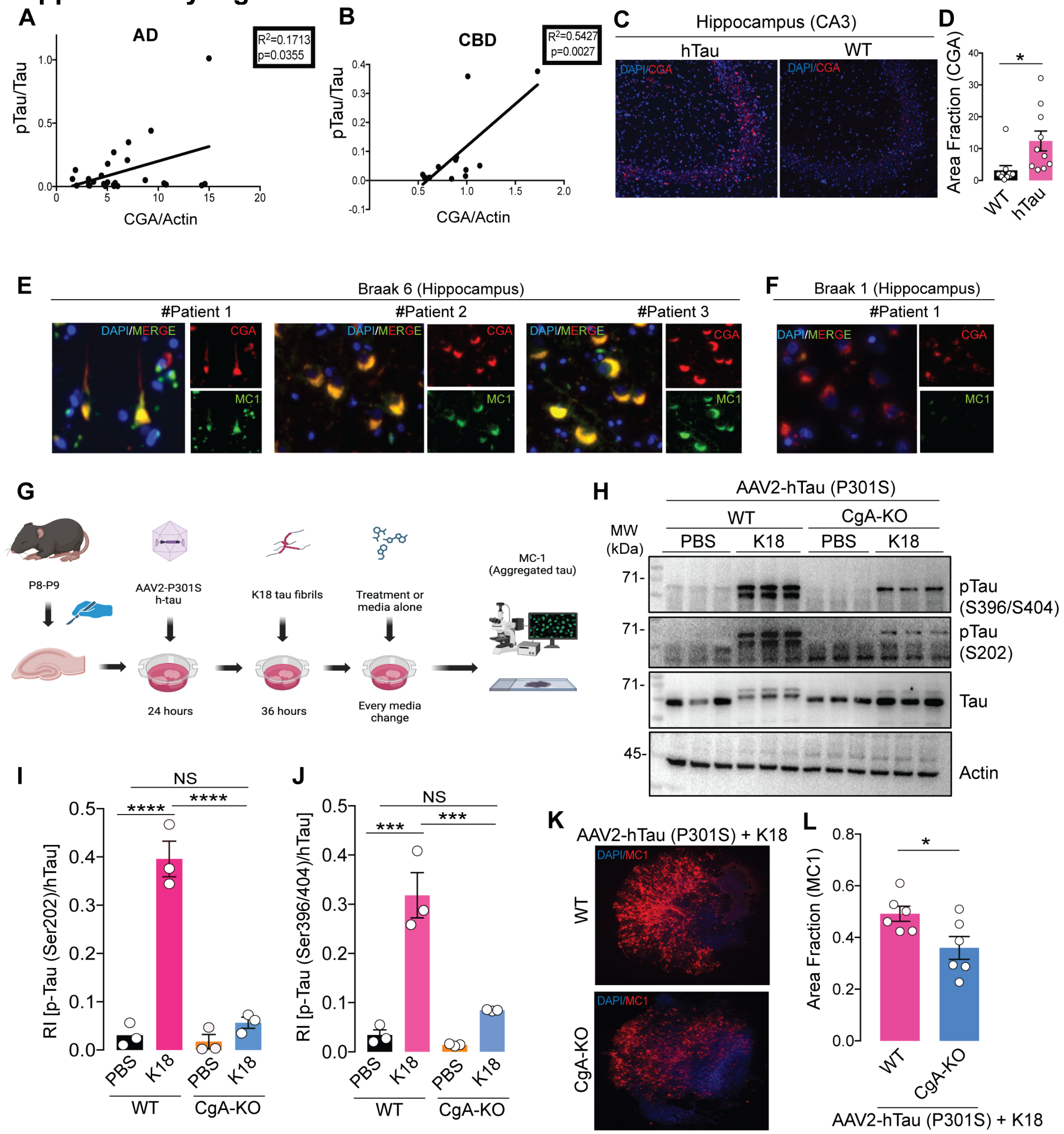

#### Supplementary Fig 2

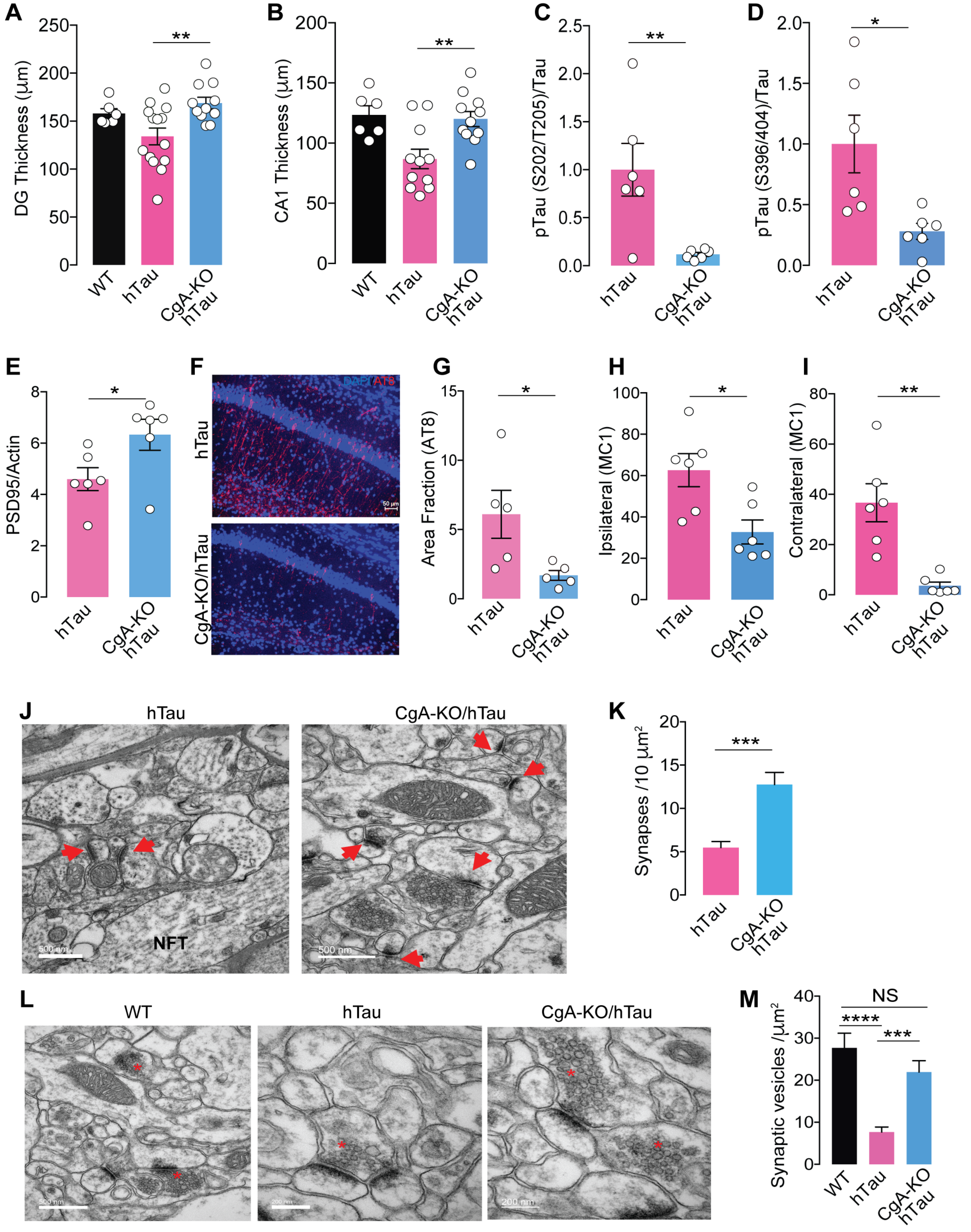

#### Supplementary Fig 3

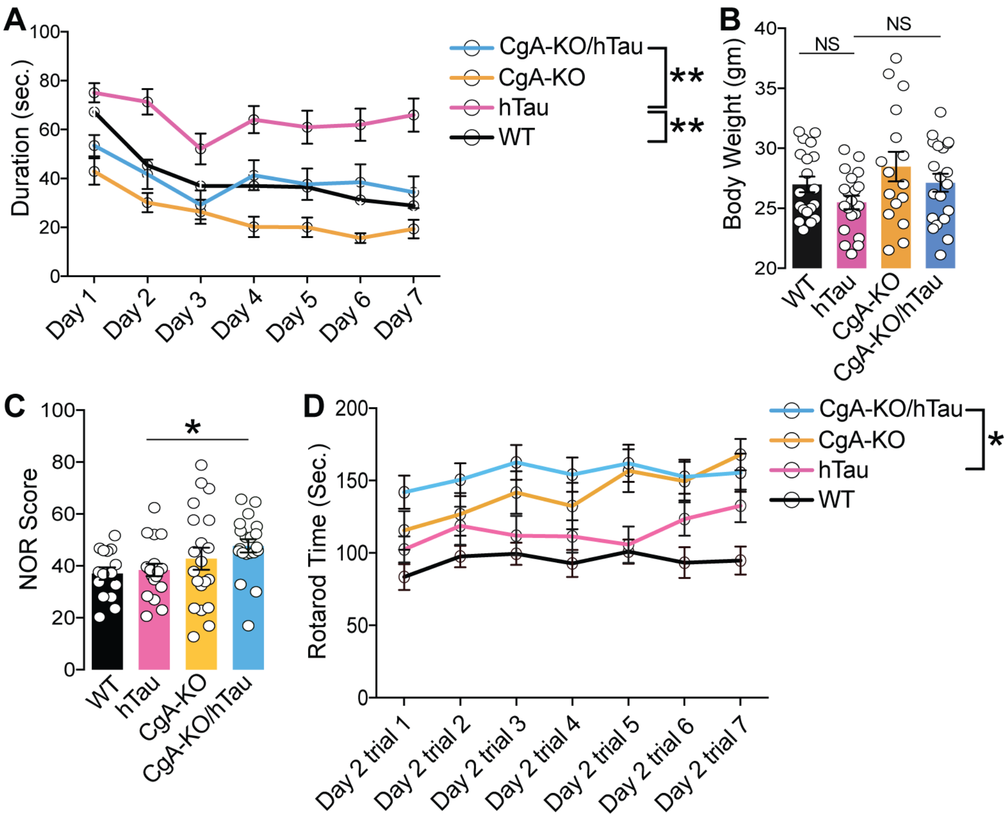

#### Supplementary Fig 4a

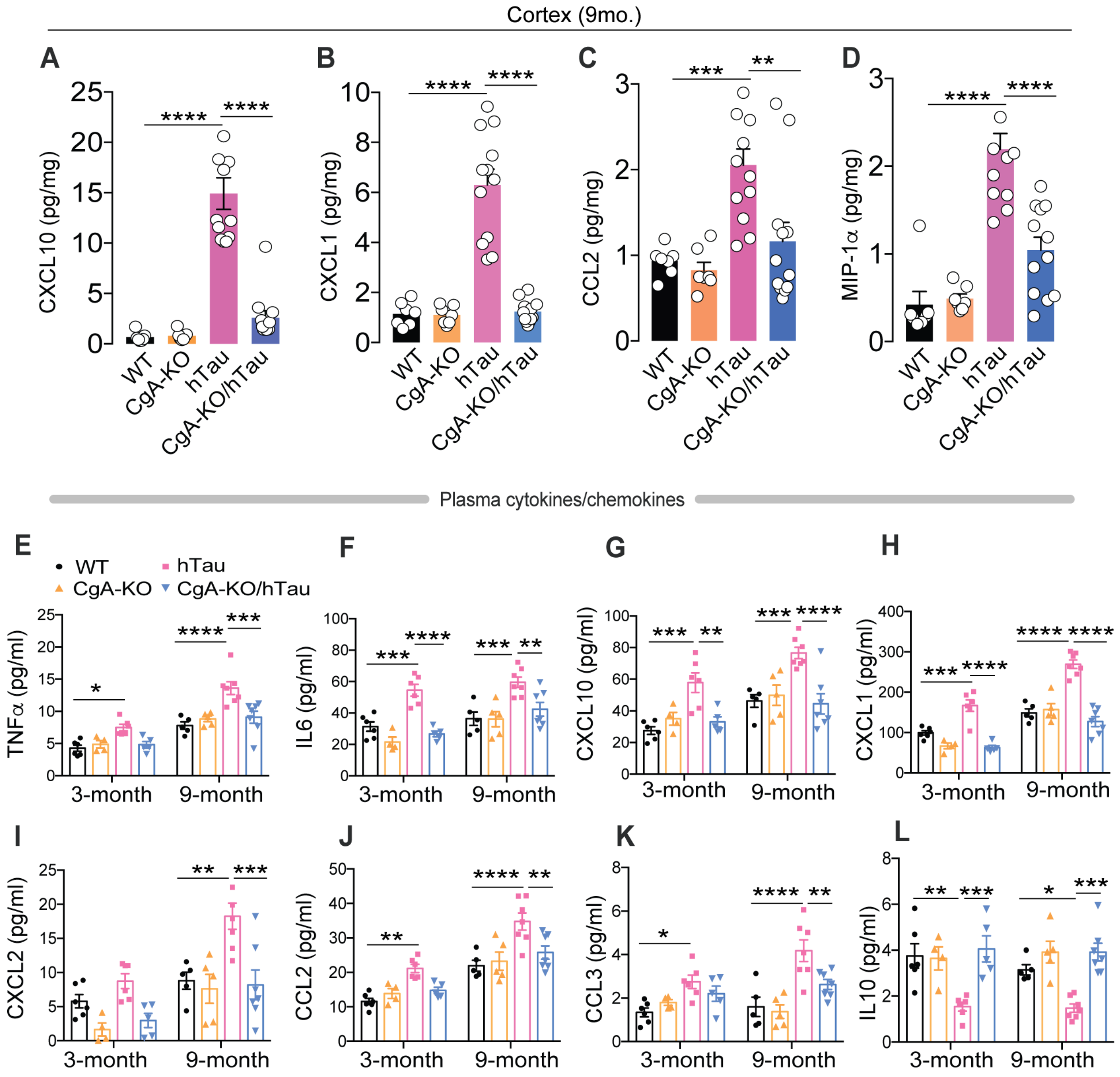

#### Supplementary Fig 4b

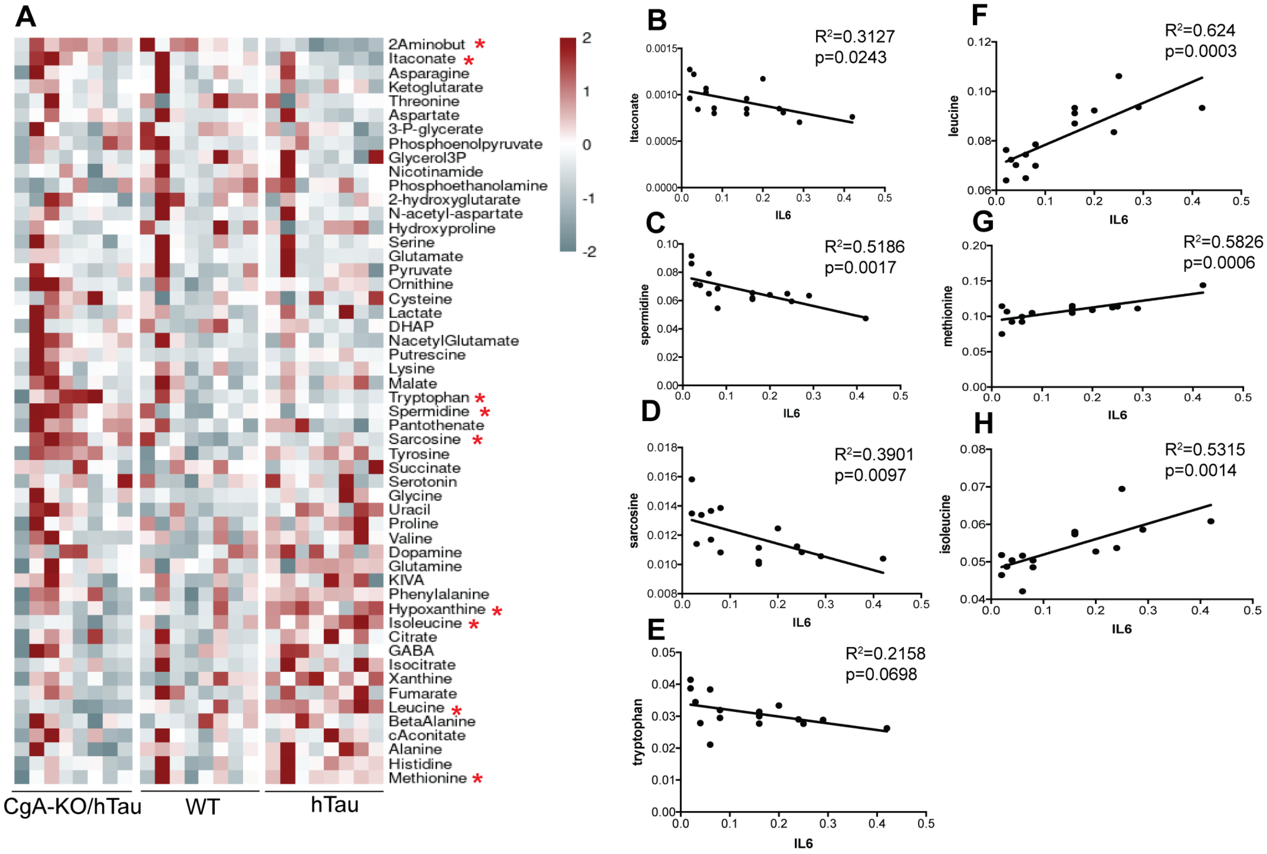

### Supplementary Fig 5

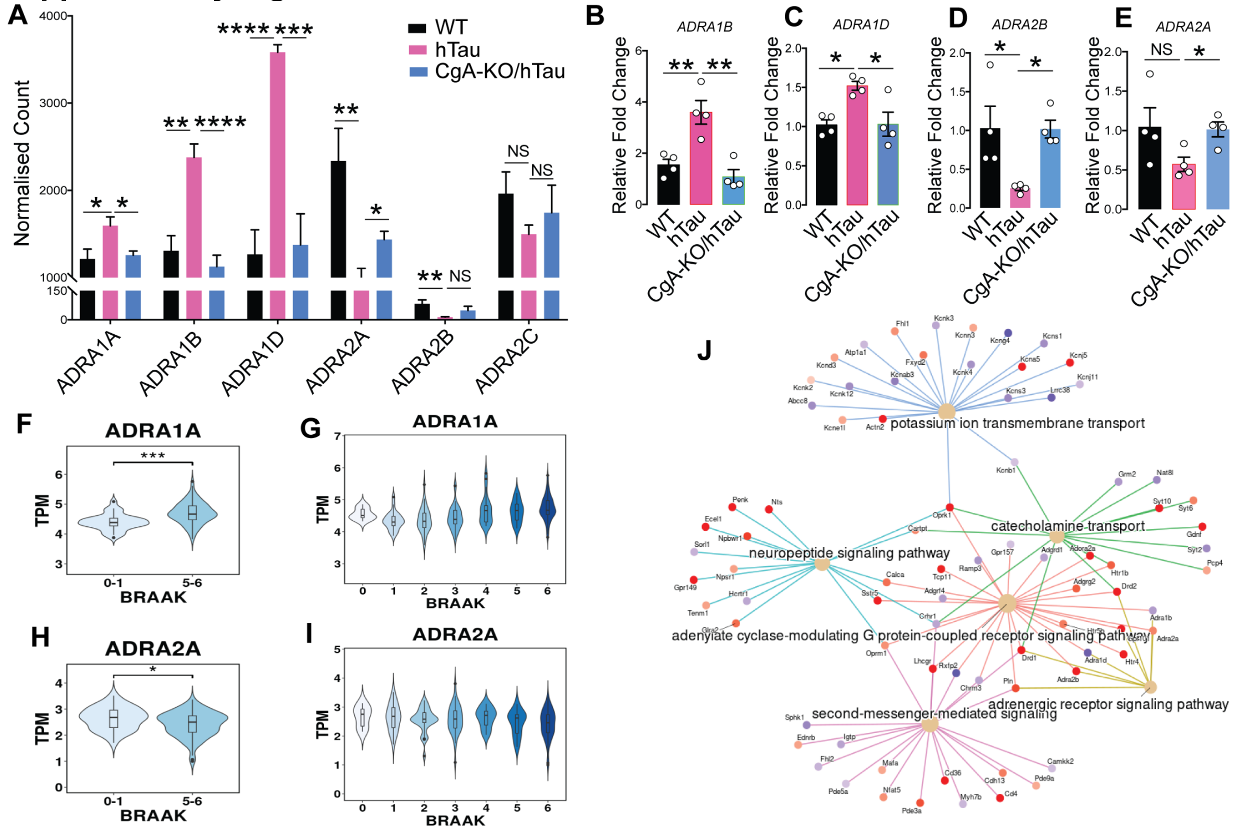

#### Supplementary Fig 6

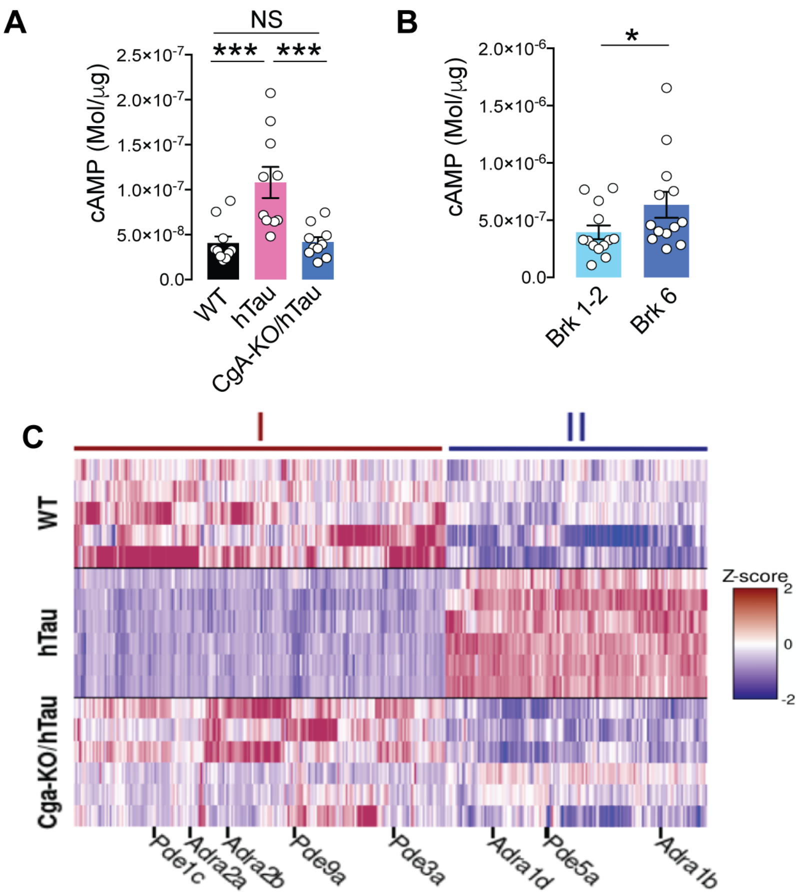

**Supplementary Fig 7**

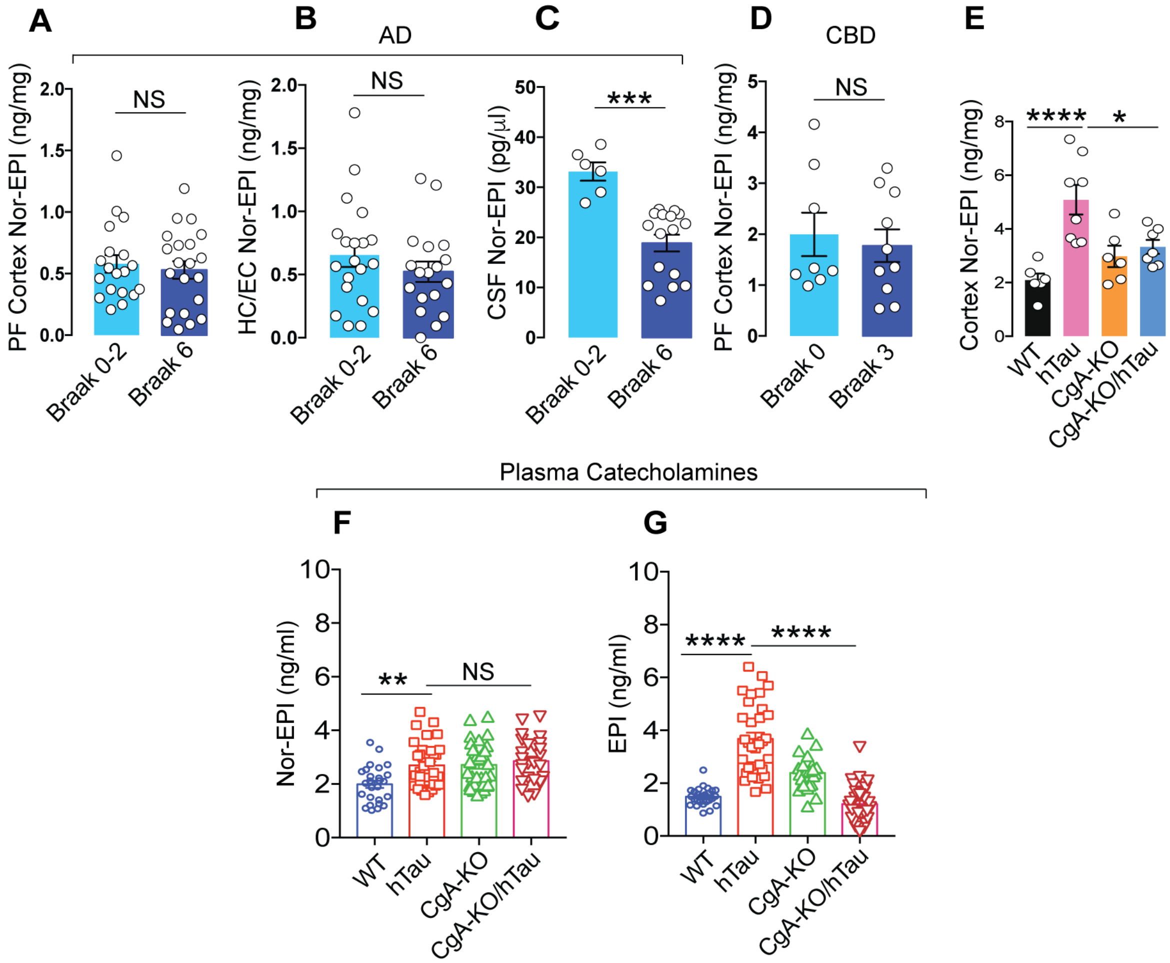
